# Dual Effect of RNA-like Polyelectrolytes on Stability and Dynamics of Biomolecular Condensates: A Tale of Competitive Short and Long-range Interactions

**DOI:** 10.64898/2026.01.29.702704

**Authors:** Deepak Sharma, Sudhriti Roy, Milan Kumar Hazra

## Abstract

Heterotypic biomolecular condensates underpin the spatiotemporal organization of cellular components and enable precise biological regulation. These condensates frequently comprise proteins and RNA at variable stoichiometries, with RNA acting as a key modulator through its ability to engage in both long-range electrostatic and short-range specific interactions. How such RNA-like components regulate condensate stability and dynamics across distinct interaction regimes, however, remains unclear. Here, we examine the role of RNA-like polyelectrolytes in tuning the stability and material properties of heterotypic condensates formed by designed short peptide sequences spanning a continuum from long-range electrostatics-to short-range hydrophobic interactions. By systematically strengthening short-range hydrophobic interactions while varying polyelectrolyte composition, we uncover a dual, regime-dependent role of RNA-like species. In electrostatics-dominated condensates, excessive polyelectrolyte concentration rapidly destabilizes droplets due to enhanced long-range repulsive interactions while the droplets retain stability until intermediate polyelectrolyte concentration fairly well. In contrast, in strongly hydrophobic condensates, polyelectrolytes function as multivalent sticker hubs, stabilizing condensates through favourable peptide– polyelectrolyte interactions. Polyelectrolyte enrichment within condensates is nonlinear with respect to mixing fraction and saturates at largest cluster mole fractions of ∼0.2–0.25. Condensate dynamics reflect this interplay: polyelectrolyte diffusivity is 20–40% higher in electrostatics dominated systems than in hydrophobic ones, while in extremely hydrophobic condensates, polyelectrolytes diffuse ∼50% more slowly than peptides, indicative of scaffold-like behaviour. Together, these results reveal tuneable and opposing roles of RNA-like polyelectrolytes in shaping condensate stability, dynamics, and morphology across diverse interaction regimes.

## Introduction

Heterotypic condensates formed by various multivalent peptides^1–4^ and RNAs, tune several cellular processes namely ribosome biogenesis^5^, stress response^6^ and even contribute toward fatal solid-like aggregation prone disease related states^7^. Several RNA-binding proteins undergo spontaneous condensate formation through demixing of proteins and nucleic acids into two distinct phases differentiated by density^8^. In all domains of life proteins and RNA based nano-to-micron scale bodies play key role in RNA synthesis, processing and degradation. For example, nucleolus, nuclear speckles and stress granules act as transcription hubs^9–13^.

Disordered proteins along with single-stranded RNA molecules undergo ordered vesicle-like assemblies at specific mixing of the components. Experiments suggest vesicular architecture is generic to multicomponent heterotypic systems undergoing phase separation. It has been speculated such vesicular assembly influences formation of liquid-like multilayered organelles that has potential to fabricate novel stimuli responsive systems^14^.

Liquid-like condensates formed by RNA binding proteins are metastable in nature^15,16^ and transforms toward a solid-like assembly over time called “aging” in a protein specific manner^17^. Such dense polymer network may undergo percolation transition^18^ in dense media, amyloid structure ^19^formation or a glass-like transition^20^ in a time irreversible manner. It has been pointed out RNAs in a sub-stoichiometric quantity in condensates act as scaffolds in enhancement of peptide-protein interaction thus acting as a core component in various RNA-protein assemblies (RNPs)^21–23^.

Non-coding RNAs like NEAT1-1 and NEAT1_2 plays a critical role in defining condensate architecture of paraspeckles^24,25,26^. Small nuclear RNAs and MALAT1 introduces structural integrity of nuclear speckles^27,28^. Protein coding mRNAs namely histone pre mRNA architect histone locus body formation^29^. Longer mRNAs form scaffolds for cytoplasmic RNP networks coordinating with fragile X-related protein 1(FXR1) and minutely regulates signalling mechanism^30^.

RNA length showcases non-trivial effect on the stability of biomolecular condensates^31^. RNA has the ability to prevent protein conformation transitions mediated alteration of viscoelasticity of the condensates^32,33^. On the other hand, RNA may also play the role of a surfactant preferably localized over the surface of a condensate modulating its size, structure and morphology^34–36^. It has been proposed that rRNA may act as surfactant to stabilize fibrillar centres of nucleolus^35^. Hence, RNA’s role in condensed phase is diverse and highly context specific in tuning size, stability and material properties^14,36–42^. Regulation of condensate’s material properties is an important aspect to study as the same may be tuned to create a spectrum of droplet states modulated by protein-RNA interactions having diverse viscoelasticity. RNA influences condensate metastability and liquid-to-solid transitions for proteins namely FUS, hnRNPA1, TDP43, TAF15, and EWSR1^43,44^. Experimental studies suggest RNA may act as heterotypic buffer in peptide-peptide interactions reducing phase separation ability of peptides^4535^. Whereas studies also have shown RNA’s potential ability to form solid amyloid-like phase transitions in-vitro and in-live cells^46,47^. Even RNAs themselves undergo phase separation with a lower critical solution temperature^48^ and may promote cellular RNP assembly formation^49^. Phase separation of Repeat expanded RNA has been shown to initiate neurological disorders namely Huntington’s disease and ALS^50–52^. The molecular mechanism and quantification is yet unclear of such dual role RNA in condensates. It has been also proposed that RNA may play role of ATP independent chaperon in condensate fusion and exchange materials, is supported by its control on SG assembly through buffering mRNA interconnects^53,54^. Alteration of peptide component or their sequence nature can influence specific organelle’s growth, fusion and even dynamics and in turn its functionality. Although RNA’s role has been pointed out in condensates, underlying molecular determinants for formation, regulation and function of RNP assemblies is yet unclear.

Cellular RNA-binding proteins has the capability of undergoing homotypic phase separation to form assemblies and constitutes an alternate regulation mode^7,16,55–64^. RNA-mediated re-entrant phase behaviour has been observed by condensation and subsequent de-condensation with enhancement of RNA enrichment^22,65–67^. Stoichiometric variation in condensate distinctly alters morphology of the same leading toward hollow vesicle like ones^14^. Such observations suggest a molecular level critical dependence of binary protein-RNA interactome on stoichiometric enrichment in dense media of both the components and not yet elucidated.

To elucidate the role of RNA-like polyelectrolytes in modulating the stability and dynamics of heterotypic condensates formed by diverse peptide sequences, we systematically varied peptide–polyelectrolyte stoichiometry in condensate-forming mixtures while monotonically tuning peptide sequences across the full interaction spectrum, from long-range electrostatic to short-range hydrophobic dominance^68,69^. The peptide interaction landscape is parameterized by the fraction of hydrophobic residues in the sequence.

Present study provides a systematic, molecular-level framework for understanding how RNA-like polyelectrolytes regulate condensate-forming propensity of intrinsically disordered peptide sequences spanning a broad spectrum of long- to short-range interactions. By quantitatively comparing condensate stability, internal dynamics, and chain reconfiguration timescales of peptides and ssRNA-like polyelectrolytes, we show how peptide–polyelectrolyte mixing fractions and sequence hydrophobicity jointly control condensate stability and dynamics. Importantly, this work establishes a well-defined baseline dominated by long-range electrostatic and short-range hydrophobic interactions, enabling the precise isolation and quantification of cation–π interaction effects in our future studies.

## Models and Simulations

### (A) Designed Systems

To investigate how a spectrum of long-to short range interactions prevailing in sequences in combination with ss-RNA like polyelectrolyte’s mixing extent alter the stability and internal dynamics of condensates, we have designed 40-residue peptide sequences with varied content of hydrophobic residues (f=0.0, 0.1, 0.2, 0.3 and 0.6) that has been used in simulations to form condensates (Fig 1). While f=0.0 denotes polyampholyte peptides with electrostatic nature, f=0.6 denote polyampholytes enriched with hydrophobic residues. We have ensured neutrality of the designed sequences to remove the effect of net-charge of sequence. For, the sake of clarity, we are only showcasing the data for four such sequences namely f=0.0, 0.1,0.3 and 0.6. A 20-residue polyelectrolyte (resembling ss-RNA) having both long and short-range interactions on each nucleotide site has been mixed systematically with peptides at distinct mole fractions (denoted as *X_polyelectrolyte_*) to introduce stoichiometry driven heterotypic condensate formation. However, we call them as “RNA like polyelectrolytes” and not exact RNA since we have not introduced the effect of cation-π interactions in the present model and only have dominant long-range electrostatics and short-range hydrophobic interactions. However, present study creates a standpoint and gives the precise control to uncover the effect of only cation-π interactions in real protein-RNA condensates being the reference and is subject to our ongoing efforts. Several mixing fractions of the peptides and polyelectrolytes has been simulated namely *X_polyelectrolyte_* = 0.05, 0.10,0.15,0.20,0.25,0.30. Representative snapshots of condensates at same distance from criticality has been shown for peptide sequences with dominant electrostatic (f=0.0) and hydrophobic short-range sites (f=0.6) at a constant disordered polyelectrolyte mixing fraction *X_polyelectrolyte_* = 0.15. While positively charged amino acids has been coloured with blue and negative ones with red, hydrophobic amino acids has been shown with limon green. Polyelectrolytes resembling RNA molecules has been shown with orange. While electrostatic interaction dominated condensates showcase a hollow droplet formation with relatively lower interactions and connected by polyelectrolytes, a dense hydrophobic condensate with polyelectrolytes being the core has been observed at intermediate polyelectrolyte concentration.

**Fig 1.**
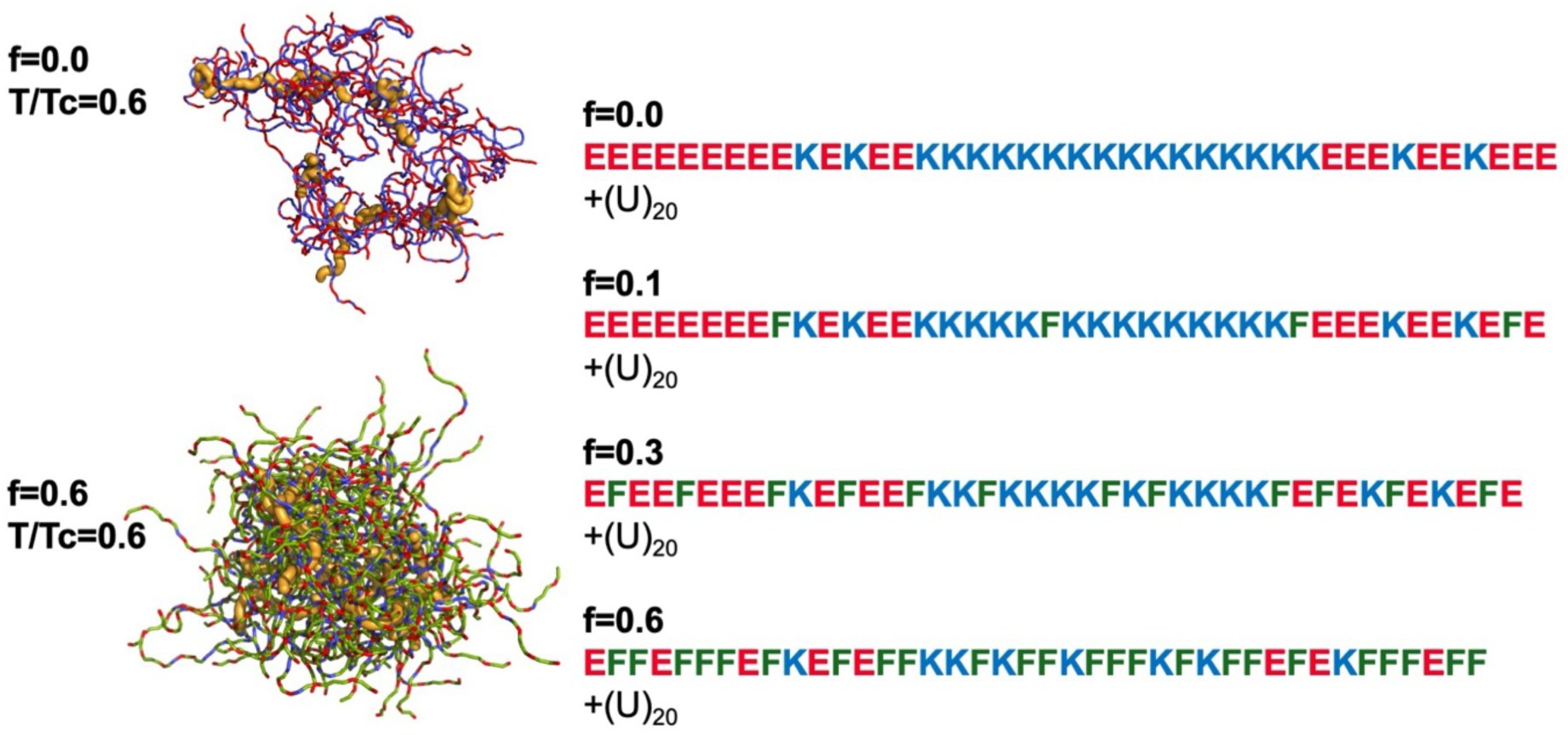
Designed peptide sequence variants with differential hydrophobicity. Heterotypic peptide-polyelectrolyte pairs have been shown with differential sequence hydrophobicity fraction f=0.0, 0.1,0.3 and 0.6. A 20-residue poly-Uracil unit act as ssRNA like polyelectolyte Representative condensates has been shown for f=0.0 and (U)_20_ and **f=0.6 and (U)_20_** pairs at *X_polyelectrolyte_* = 0.15 and T/Tc =0.6. Positively charged amino acids has been shown with blue and negative ones with red, hydrophobic amino acids has been coloured with limon green. Polyelectrolytes resembling RNA molecules has been shown with orange.

### Simulation Details

Because all-atom explicit-solvent simulations are too computationally expensive to reach the timescales required for equilibrium mesoscopic condensate formation, we employ a coarse-grained model^68,69^ to study equilibrated heterotypic condensates and systematically dissect how heterogeneity influences their stability and dynamics.

Each residue in the designed polyampholytes/polyelectrolytes has been represented by a single bead and assigned either a positive charge (for Lys), negative charge (GLU and Uracil of polyelectrolytes) and charge neutrality for hydrophobic phenyl alanine sites (PHE). The potential energy function includes harmonic bond and angular terms defining the polymer architecture, electrostatic interactions among all charged beads, and short-range dispersion interactions. Dihedral interactions are omitted, as the designed peptides and polyelectrolytes behave as disordered entities having ensemble of conformations.

An embedded implicit solvent model and salt effect has been introduced to screen electrostatic interactions among charged residues through Debye-Hückel potential^70^.

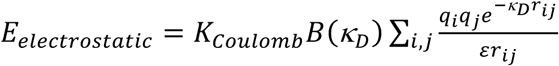

where 𝑞_𝑖_ and 𝑞_j_ denote the charge of the *i*^th^ and j^th^ bead, 𝑟_𝑖j_ denotes the inter-bead distance, 𝜀 is the solvent dielectric constant, and 𝐾_𝐶𝑜𝑢𝑙𝑜𝑚𝑏_ = 4𝜋𝜀_O_ =332 kcal/mol. 𝐵(κ_𝐷_) is a function of solvent salt concentration and the radius (𝑎) of ions produced by the dissociation of the salt, an can be expressed as

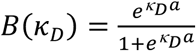

The Debye-Hückel electrostatic interactions of an ion pair act over a length scale of the order of 𝜅^−1^, which is called the Debye screening length. κ_𝐷_ is related to ionic strength as,

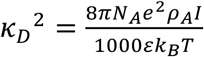

where 𝑁_A_is the Avogadro number, *e* is the charge of an electron, 𝜌_A_ is the solvent density, *I* denotes solvent ionic strength, 𝑘_B_is the Boltzmann constant, and T is the temperature. To avoid overlap among beads, a steep repulsion interaction has been defined as

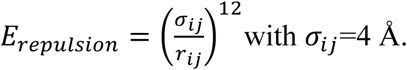

In addition, hydrophobic residues interact with a short-range 12-10 dispersion interaction

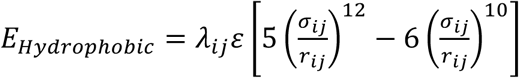

In our simplest coarse-grained representation, we have implemented Phenyl-Alanine-Phenyl Alanine (PHE-PHE) and Phenyl Alanine-Uracil (PHE-U) short range interactions respectively. We have used 𝜀 = 0.2 𝑘𝑐𝑎𝑙/𝑚𝑜𝑙 that predicts effective dimension of disordered peptides similar to experiments^68,71^ and scaling parameter 𝜆_𝑃𝐻𝐸_ = 1.0 as described in HPS model^71,72^ for extremely hydrophobic PHE. Uracil being very similar to Thymine in structural perspective, we have used scaling parametrization for thymine with an enhanced contribution from methyl-group of Uracil as 𝜆_𝑁𝑢𝑐𝑙𝑒𝑜𝑡𝑖𝑑𝑒_ = 0.92 cumulating the contribution from poly-U backbone and base hydrophobicity contribution in a single nucleotide bead. VDW radius of PHE residue and a nucleotide bead is comparable (∼3.5 Å), hence the diameter (𝜎_𝑖j_) has been set to 7 Angstrom for both pairs of short-range contacts.

Langevin dynamics simulations were performed for 100 copies of binary polymer pairs at varying mixing fractions (*X_polyelectrolyte_*) in a cubic box of size 300 × 300 × 300 Å³. To characterize single-chain properties in the dilute phase, individual polymers were simulated in the same box at temperatures below the critical point of the corresponding condensates. Condensate stability and polymer dynamics were analysed at an ionic strength of 0.02 M using an implicit solvent with a dielectric constant of 80, across multiple subcritical temperatures.

At each temperature, two independent trajectories of 10⁷ steps were generated by integrating the Langevin equation to ensure convergence and statistical averaging. Configurations were saved every 500 steps, yielding 20,000 frames per trajectory. The first 5,000 frames (2.5 × 10⁶ steps) were discarded for equilibration, and the remaining 15,000 frames were used for analysis. Condensates were identified using a clustering algorithm, and the largest clusters were selected for subsequent analysis.

## Results and Discussion

### Dual role of RNA-like polyelectrolytes in altering stability of condensates

To understand how short single-stranded RNA-like negatively charged polyelectrolytes alter the phase behaviour of designed sequences ranging from electrostatic to hydrophobic in nature (Fig 1), temperature-peptide fraction phase diagrams has been plotted for systems composed of sequences having considerably differential hydrophobicity and at different polyelectrolyte mixing fractions (Fig 2). At each timestep we have computed the fraction of total polymers in largest cluster and in dilute phase. A Caylee Tree like clustering algorithm has been employed to define the largest cluster. The critical point has been determined by fitting polymer fraction-temperature data to the Ising like model with the following expression

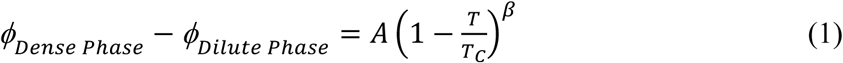

**Fig 2.**
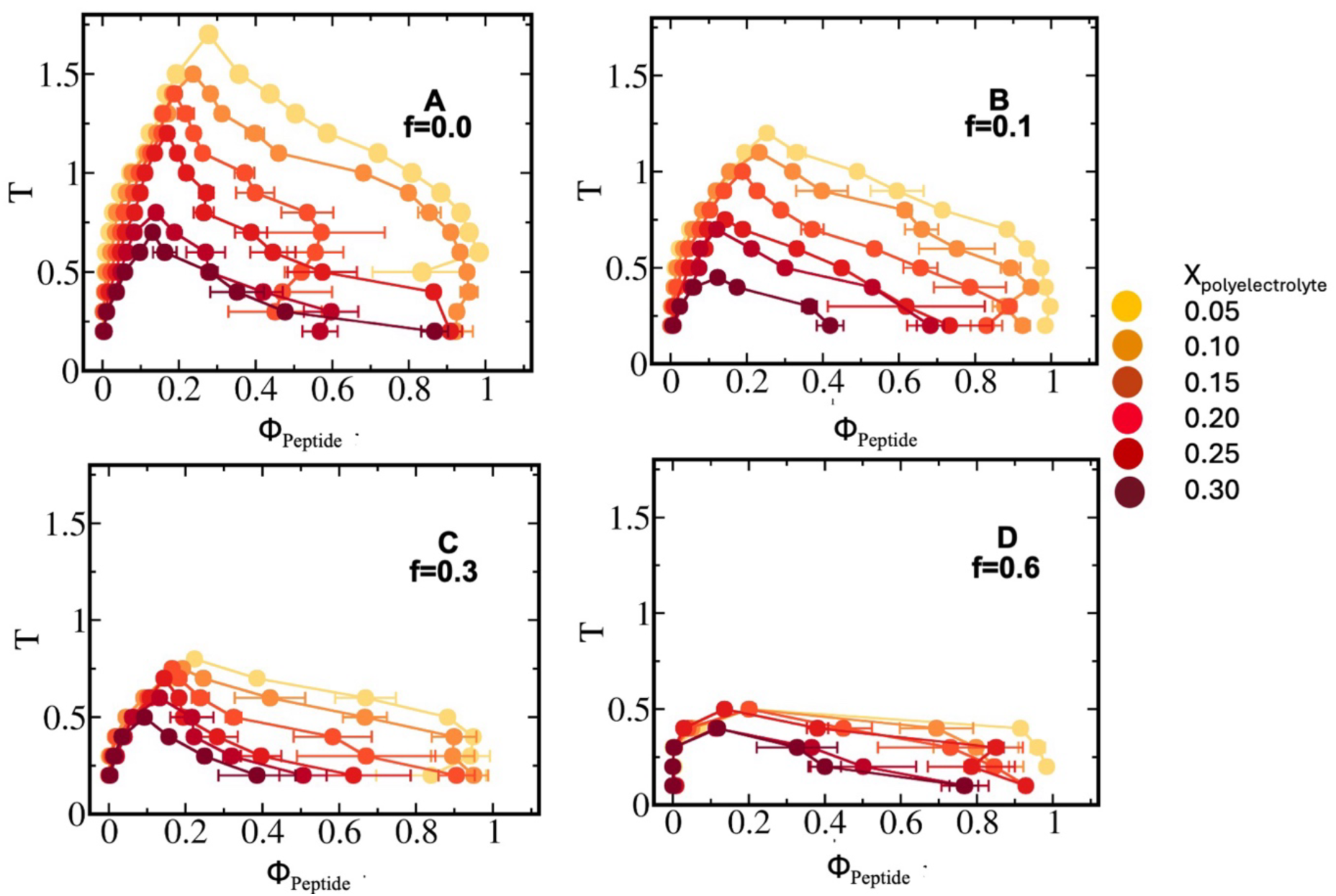
Phase diagram and stability of the condensates formed by studied heterotypic pairs. (A)f=0.0 and (U)_20_, (B) f=0.1 and (U)_20_ and (C) f=0.3 and (U)_20_ (D) f=0.6 and (U)_20_ Different mixing fractions has been shown in each panel ranging from *X_polyelectrolyte_* = 0.05 − 0.30 with yellow-dark red color.

While designed peptides having significantly less hydrophobicity (denoted as f, Fig 1) show extensive destabilization of condensate phase at excessive concentration of polyelectrolytes retains its stability until intermediate polyelectrolyte enrichment. The effect diminishes as sequence hydrophobicity enhances.

Condensates formed by sequences having hydrophobicity fraction f=0.0, 0.1 and 0.3 showcase nearly 4-1.5-fold decrease of stability respectively with enhancement of polyelectrolytes (Fig 2 panels A-C). Condensates formed by peptides with significant hydrophobicity (f=0.6) showcase almost no effect of polyelectrolyte enrichment (Fig 2 panel D).

One must note each residue in polyelectrolyte can form hydrophobic attractive short-range contacts with peptide hydrophobic sites in addition to salt-bridges among charged residues. Once the hydrophobicity of sequences forming condensates enhance, we observe electrostatic repulsion among like-charges of peptide-polyelectrolyte pairs being replaced by short-range attractive stickers that can connect more peptides as a scaffold and leading to unaltered stability of the condensate.

### Underlying energetics of phase stability and connection to criticality

To decode the origin of such alteration of stability along polyelectrolyte mixing fraction, total energetic stabilization of the condensate polymers with respect to isolated dilute phase ones has been shown at different distances from criticality and at various polyelectrolyte mixing fractions (**Fig 3**). In addition, total energy has been dissected into components namely self-interactions among peptides, polyelectrolytes and cross interactions among the two kinds (Fig S1-4).

**Fig 3.**
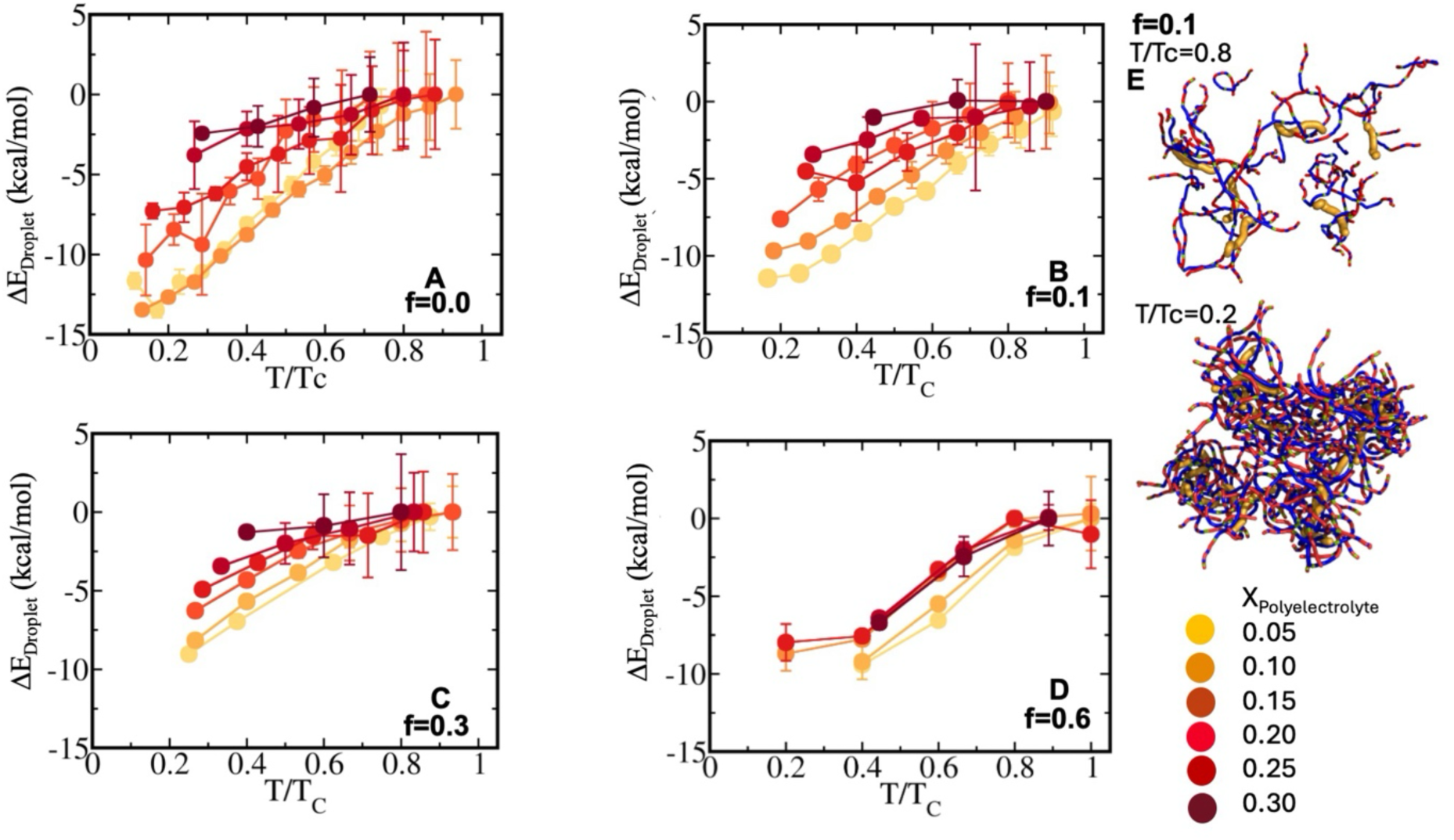
Energetic stabilization of peptides in condensate phase with respect to bulk. Relative energetic stabilization, each polymer faces in condensate phase with respect to dilute ones for peptide-polyelectrolyte sequence pairs with (A) f=0.0 and (U)_20_, (B) f=0.1 and (U)_20_, (C) f=0.3 and (U)_20_ (D) f=0.6 and (U)_20_ Different mixing fractions has been shown in each panel ranging from *X_polyelectrolyte_* = 0.05 − 0.30 with yellow to dark red color. (E) Representative snapshots of peptide-polyelectrolyte network has been shown for f=0.1 and (U)_20_ at near (T/T_C_=0.8) and far away (T/T_C_=0.2) from critical point. Owing to the dominance of long-range interactions in these sequences, peptide–polyelectrolyte small clusters and even oligomeric assemblies persist near and beyond criticality. Total energy, a peptide experiences in droplet phase is composed of self and cross components. Individual components namely peptide-peptide (P-P), polyelectrolyte-polyelectrolyte (PE-PE) self-interaction and peptide-polyelectrolyte (P-PE) cross ones has been shown in Fig. S1-4 for designed peptide sequences.

For sequence having completely electrostatics nature (f=0.0), energetic stabilization of condensate phase polymers (ΔE_Droplet_) with respect to dilute phase stemming from electrostatic interactions among peptide-peptide and peptide-polyelectrolyte ones, contribute around 10 kcal/mol at X_Polyelectrolyte_ =0.05 and decays monotonically to 2.5 kcal/mol upon variation of X_Polyelectrolyte_ at T/Tc =0.4 signifying nearly a 4-fold loss of energetic stability (Fig 3A).

For sequences with low-to-moderate hydrophobic content (f=0.1 and 0.3 panel 3B and C), condensate phase at X_Polyelectrolyte_ =0.05 and T/Tc=0.4 has relatively lower stability than condensate formed by f=0.0 sequence associated with energetic gain of nearly 7-8 kcal/mol. A prominent gradual decrease in stability has been observed (as in Fig 1B and C) and energetic loss is nearly 7-fold upon variation of X_Polyelectrolyte_ (Fig 3B).

At extreme hydrophobicity of designed sequences as peptides interact with RNA like polyelectrolytes with majorly short-range hydrophobic interactions (f=0.6, Fig 3D), the loss of energetic stabilization due to variation of polyelectrolyte mixing fraction is only 3 kcal/mol at T/Tc=0.4 and much lesser than condensates formed by long-range electrostatic interactions.

As temperature increases, droplet tends to lose stability owing to the loss of peptide-peptide interactions (Fig S1-4 panel A). Surprisingly peptide-polyelectrolyte interactions showcase nearly no notable temperature dependence (Fig S1-4 panel B) that points toward an oligomeric state of peptides and polyelectrolytes may always persist even at high temperature. For f=0.1, at X_Polyelectrolyte_ =0.15, snapshots reveal a dense polymer–peptide mesh at T/Tc=0.2 that transforms into a void-rich, two-dimensional network near criticality T/Tc=0.8, electrostatically mediated by polyelectrolytes (Fig. 3E). Peptide-polyelectrolyte cross interactions contribute nearly 5 kcal/mol stabilization at X_Polyelectrolyte_ =0.05 and nearly 25-30 kcal/mol stabilization at X_Polyelectrolyte_ =0.30 and T/Tc=0.4 indicating a 5 to 6 times enhancement upon variation of sequence hydrophobicity. Repulsive self-interactions among polyelectrolytes diminish with enhancement of temperature below criticality as condensates break apart into smaller ones (Fig S1-4 panel C).

To decode further into individual pairwise energetics, we have observed polyelectrolyte concentration affects peptide-peptide self-interactions significantly due to dominant repulsive long-range electrostatics and primarily due to loss of peptide component that could buffer the repulsions in the system (Fig S1-4 panel A). For f=0.0 sequence, a peptide in condensate phase experiences nearly 9 kcal/mol energetic stability at X_Polyelectrolyte_ =0.05 and at T/Tc=0.4 that diminishes to 0.5 kcal/mol at X_Polyelectrolyte_ =0.30 (Fig S1 panel A). One may conclude the primary component of droplet stabilization is self-interactions among peptides while peptide-polyelectrolyte pairs act as core interactions for inherent stability of droplets (Fig S1-4). At moderate hydrophobicity content in sequences (f=0.1 and 0.3), peptide-peptide self-interactions lose nearly 13-15 kcal/mol stability for each peptide in condensate while varying polyelectrolyte mixing fraction from X_Polyelectrolyte_ =0.05 to 0.30 at a constant distance from criticality T/Tc=0.4 (Fig S2 and 3 panel A). At extreme limit of hydrophobicity in sequences (f=0.6) the loss of peptide-peptide self-interaction is lesser and nearly 7 kcal/mol while polyelectrolyte mixing fraction is maximum which demonstrate polyelectrolytes act as stickers in stitching hydrophobic condensates (Fig S4 panel A). It is important to note the loss of self-interactions among peptides are higher for low and moderately hydrophobic sequences (namely f=0.1 and f=0.3) and lesser at the two limits where extreme electrostatic and hydrophobic interactions play dominant role indicating a competitiveness among short and long-range interactions (Fig S1-4A).

Correlation of critical point and energetic stabilization in dense phase at a fixed distance from criticality (T/T_C_=0.4) has been shown as a function of polyelectrolyte mixing fraction X_Polyelectrolyte._ (Fig 4A and B). For peptide sequences with f=0.0 and 0.1, stability of condensate phase showcase prominent non-linearity along polyelectrolyte mixing fraction, critical point of the condensates (T_C_) decreases at a slower rate until X_Polyelectrolyte_=0.15, thereafter the decrease in T_C_ is rapid. Energetic stabilization along polyelectrolyte mixing fraction showcase two distinct states of condensate stability. Below polyelectrolyte mixing fraction X_Polyelectrolyte_=0.15, condensates dominated by electrostatic interactions retains stability whereas beyond X_Polyelectrolyte_=0.15, we have observed rapid loss of energetic stabilization of droplets (Fig 4B). Droplet stability approaches a monotonic and slower decay along X_Polyelectrolyte_ as hydrophobic residue fraction (f) of the peptide sequences enhances indicating a stabilization of dense phase due to peptide-polyelectrolyte short-range crosstalk acting as hub of stickers in maximizing peptide-peptide proximity leading toward almost similar stability independent of polyelectrolyte fractions (Fig 4A and B).

**Fig 4.**
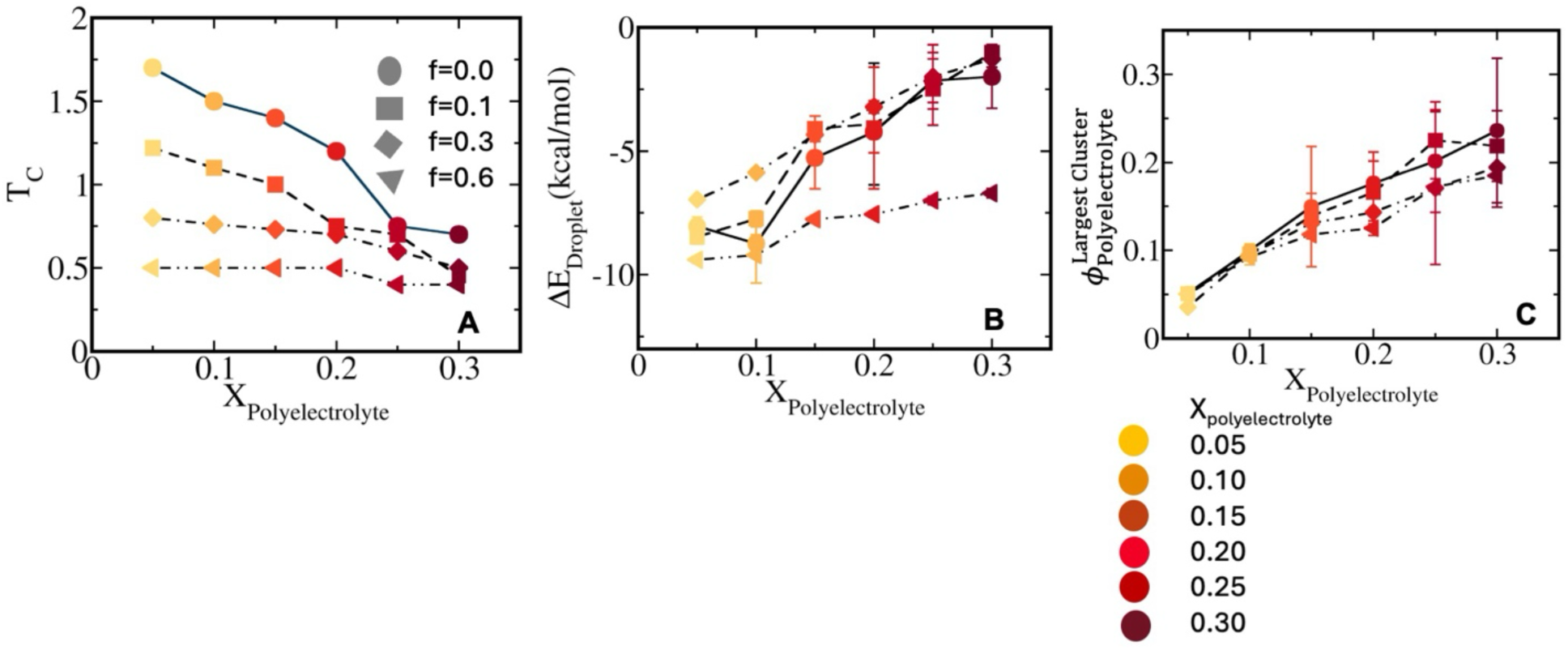
Correlated Stability-energetics and enrichment in droplets. (A)Non-linear dependence of critical point along mixing fraction *X_polyelectrolyte_* has been shown for condensate forming pairs f=0.0 and (U)_20_ (shown with circle connected with solid line), f=0.1 and (U)_20_ (shown with squares connected with dashed line), f=0.3 and (U)_20_ (shown with diamond shapes connected with dot dash line) f=0.6 and (U)_20_ (shown with triangles connected with double dot dash line). Color codes represent polyelectrolyte mixing fractions. (B) Relative energetic stabilization of peptides in condensate phase with respect to dilute phase at constant T/Tc =0.4 along mixing fraction *X_polyelectrolyte_* for all the designed peptide-polyelectrolyte variant pairs. (C) Polyelectrolyte enrichment in largest cluster (*ϕ_polyelectrolyte_*) along mixing fraction *X_polyelectrolyte_* for the studied systems shows non-linear saturated enrichment of polyelectrolytes in the designed condensates and even more pronounced with increase in peptide hydrophobicity at moderate to high *X_polyelectrolyte_*.

In addition, relative enrichment of polyelectrolytes in largest cluster has been plotted in Fig 4C. 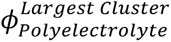 reflects the relative composition of polyelectrolytes in largest cluster. With increase in polyelectrolyte mixing fraction (X_Polyelectrolyte_) in the designed systems, we have observed a non-linear enrichment of polyelectrolytes in largest cluster across various sequence hydrophobicity. While at X_Polyelectrolyte_=0.05-0.15, relative mole-fraction of polyelectrolytes in largest cluster is nearly similar to the polyelectrolyte concentration in the system but in X_Polyelectrolyte_=0.15-0.30, enrichment in largest cluster 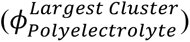 is lower than polyelectrolyte mixing fraction (X_Polyelectrolyte_) and even more so as hydrophobicity of the sequence increases which points toward a saturation of polyelectrolyte fusion in condensates i.e: lower number of polyelectrolytes relative to available polyelectrolytes acting as stickers with multiple short-range hydrophobic interaction sites and being shared by multiple peptides.

### Differential dynamics in condensate phase for polyelectrolyte and peptides

Correlated to non-linear enrichment, dynamics of components namely peptides and polyelectrolytes in condensate phase is distinctly different. To quantify the internal dynamics of polymers (namely peptides and polyelectrolytes) in droplets, we have computed mean squared displacement of peptides and polyelectrolytes at X_Polyelectrolyte_=0.15 for a constant distance from criticality T/Tc=0.6 for all the sequence variants defined as follows (Fig 5)

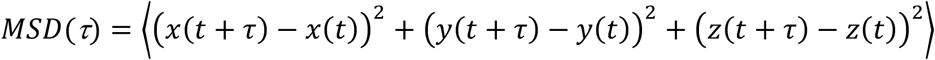

**Fig 5.**
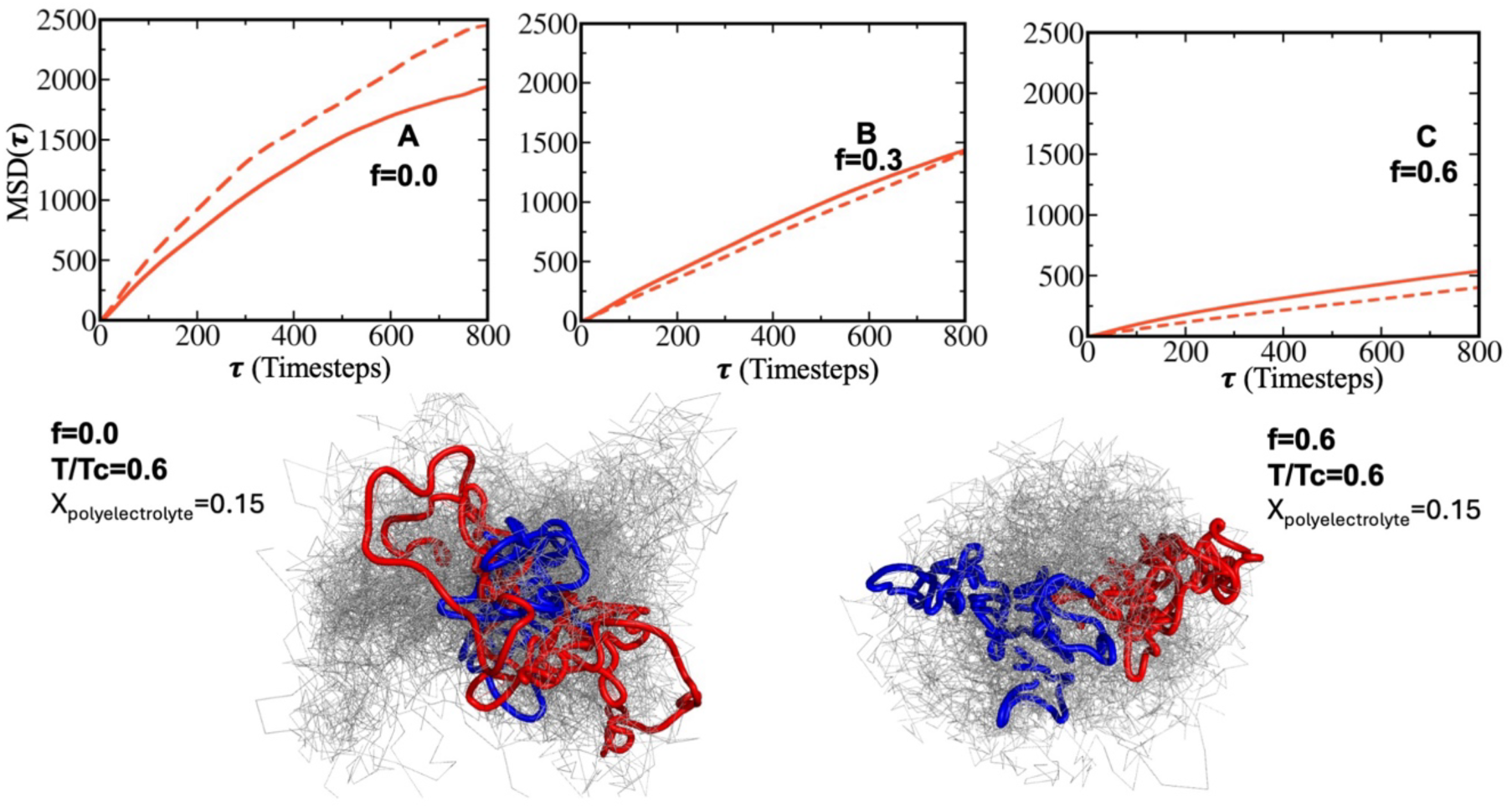
Translational diffusivity of peptides and polyelectrolytes in condensate phase. Mean squared displacement (MSD) of peptides has been shown for sequence pairs with (A) f=0.0 and (U)_20_, (B) f=0.3 and (U)_20_ and (C) f=0.6 and (U)_20_ at an intermediate mixing fraction *X_polyelectrolyte_* = 0.15 and at T/Tc=0.6. While solid line indicates MSD for peptides, dashed line indicates MSD for polyelectrolytes. Representative trajectories in condensate phase for peptides has been shown with blue and for polyelectrolytes with red in all polymer’s trajectories (grey) inside condensates formed by respectively pairs f=0.0 and (U)_20._ and f=0.6 and (U)_20_.

Convergence of mean squared displacements of peptides (Fig S5) and polyelectrolytes (Fig S6) has been shown from two independent simulations of condensate phase. In low hydrophobicity limit of sequences, we have observed polyelectrolyte’s mean-squared displacement is significantly higher than peptides in condensate phase (f=0.0, Fig 5A).

Once sequence hydrophobicity enhances, although overall diffusivity of the condensate decreases enormously, peptides showcase a relatively higher diffusivity than polyelectrolytes (Fig 5B and C). Such observation is linked to a continuous shift from long-range electrostatics interactions (being prevalent in lower hydrophobicity limit) toward a short-range interaction at higher hydrophobic sequences with polyelectrolytes being the hub of such short-range stickers (f=0.3, in Fig 5B and f=0.6 in Fig 5C). Representative trajectories has been shown for peptide (blue) and polyelectrolytes (red) among all polymers’ trajectories (Grey) at T/Tc=0.6 in condensate phase for sequences with minimum (f=0.0) and extreme hydrophobic content (f=0.6) (Fig 5D).

Ratio of average diffusivity of polyelectrolytes and peptides in condensates 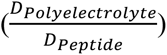 been plotted at different distance from critical point for all the designed sequences in Fig 6. Diffusivity has been calculated from mean squared displacement as 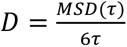.

**Fig 6.**
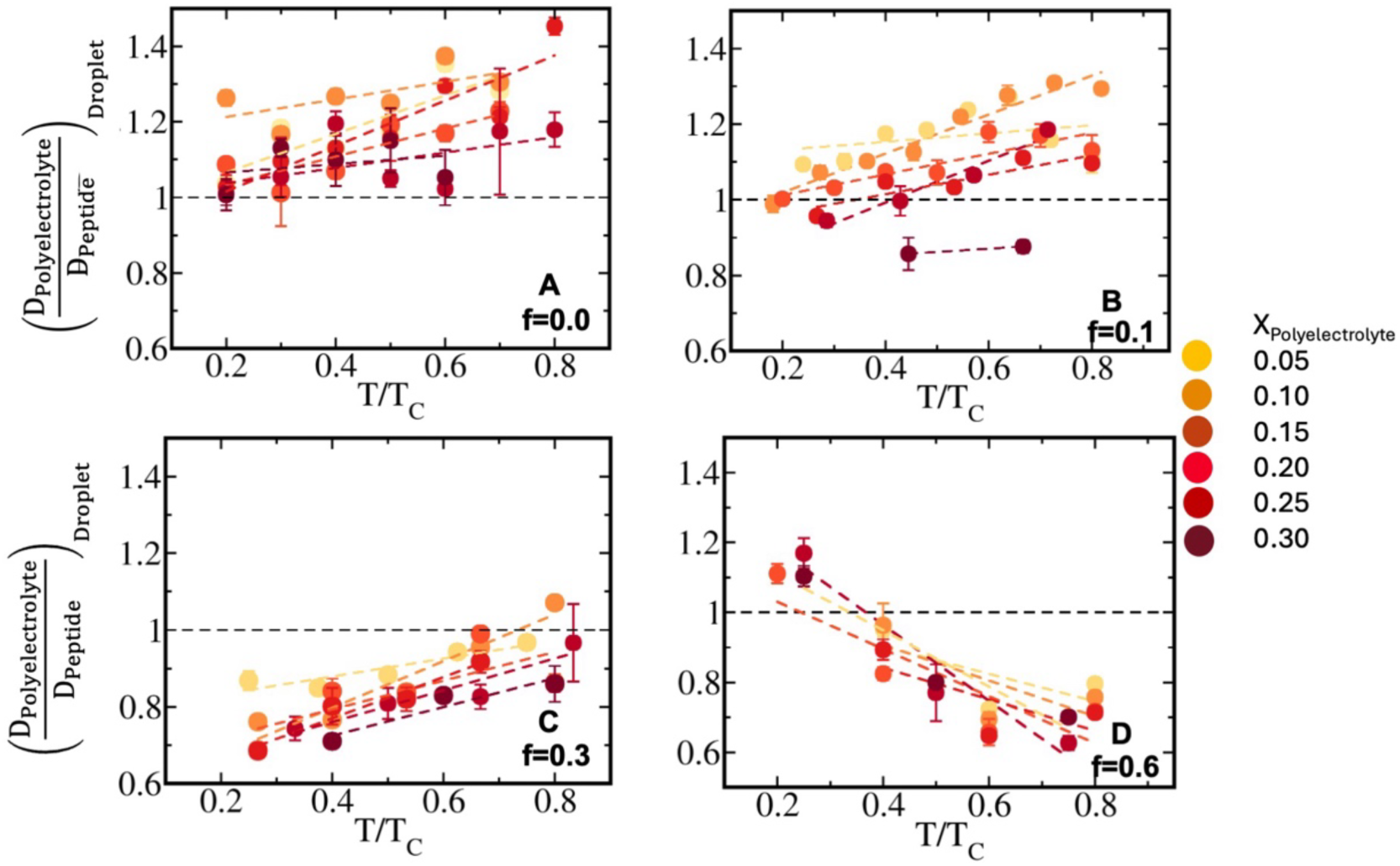
Differential dynamics of peptides and polyelectrolytes in condensate phase. Ratio of average diffusivity between polyelectrolytes and peptides 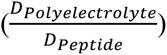 in condensate phase along temperature scaled to criticality for sequence pairs (A) f=0.0 and (U)_20_, (B) f=0.1 and (U)_20_ and (C) f=0.3 and (U)_20_ (D) f=0.6 and (U)_20_. Different mixing fractions has been shown in each panel ranging from *X_polyelectrolyte_* == 0.05 to 0.30 with yellow to dark red. At dominant electrostatic limit of peptide sequences, polyelectrolytes translates faster in condensate than peptides while at enhanced hydrophobic limit peptides are relatively faster than polyelectrolytes.

At low hydrophobic content in the designed sequences (f=0.0-0.1), polyelectrolytes diffuse within the condensate phase significantly faster than peptides, with 𝐷_Polyelectrolyte_ being ∼1.2– 1.4-times higher than 𝐷_Peptide_ at low polyelectrolyte mixing fractions. This mobility contrast further increases as the temperature approaches the critical point. As the polyelectrolyte enrichment increases, the condensates weaken and fragment into multiple smaller droplets (Fig. 1A). Concomitantly, peptide diffusivity increases and approaches that of the polyelectrolytes, causing the diffusivity ratio to decrease toward unity.

Ideally, polyelectrolytes should be twice faster than peptides due to mass scaling in absence of interactions. However, the ratio observed in condensate phase having a network of interactions are low and at most reaches 1.4. At extreme polyelectrolyte concentration (X_Polyelectrolyte_=0.30) for f=0.1 sequence, 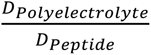 reaches to 0.85 signifying the first appearance of polyelectrolyte diffusivity being slower than peptides.

At moderately higher sequence hydrophobicity (f=0.3), polyelectrolytes exhibit a pronounced translational slowdown within the condensate relative to peptides, in contrast to electrostatically dominated sequences. This behaviour indicates that polyelectrolytes act as interaction hubs, binding multiple peptides via short-range contacts and effectively scaffolding peptide assembly (Fig. 6C vs. Fig. 6A,B). The slowdown is strongest at high polyelectrolyte enrichment (𝑋_Polyelectrolyte_ = 0.30), where polyelectrolyte diffusivity is ∼0.6 times that of peptides at T/T_C_=0.4, increasing gradually with temperature. At lower enrichment (𝑋_Polyelectrolyte_ = 0.05), polyelectrolytes remain ∼0.8 times slower than peptides at the same distance from criticality, with diffusivities converging near T_C_.

For strongly hydrophobic sequences (f=0.6), polyelectrolyte slowdown dominates and is further amplified with increasing temperature (Fig. 5C, Fig. 6D). Notably, the maximum slowdown occurs near criticality, where polyelectrolyte diffusivity drops to ∼0.6 times that of peptides, whereas at lower temperatures the diffusivities are comparable, with polyelectrolytes only ∼0.8 times slower.

We further observe a clear correlation between polyelectrolyte fraction in largest cluster and component diffusivity within the condensate. As shown in Fig. 4C, at increased peptide hydrophobicity the relative enrichment of polyelectrolytes is reduced compared to electrostatically dominated systems, at moderate to high 𝑋_Polyelectrolyte_. In this regime, a similar number of polyelectrolytes (with respect to available ones in the system) act as hubs, engaging multiple peptides through combined long- and short-range interactions to scaffold and stabilize the droplet phase. This hub-like role anchors the condensate structure, resulting in reduced polyelectrolyte diffusivity at higher hydrophobicity (Fig. 4C).

### Liquid-like nature of droplets and reconfiguration lifetimes of peptides in condensate phase

To decode the liquid-like nature of condensate phase, ratio of diffusivity of peptides in condensate phase and bulk has been plotted (Fig. 7). Comparative mean squared displacements of peptides has been shown in droplet and bulk in Fig S7 for all the designed sequences at X_Polyelectrolyte_=0.15 and T/T_C_=0.6.

**Fig 7.**
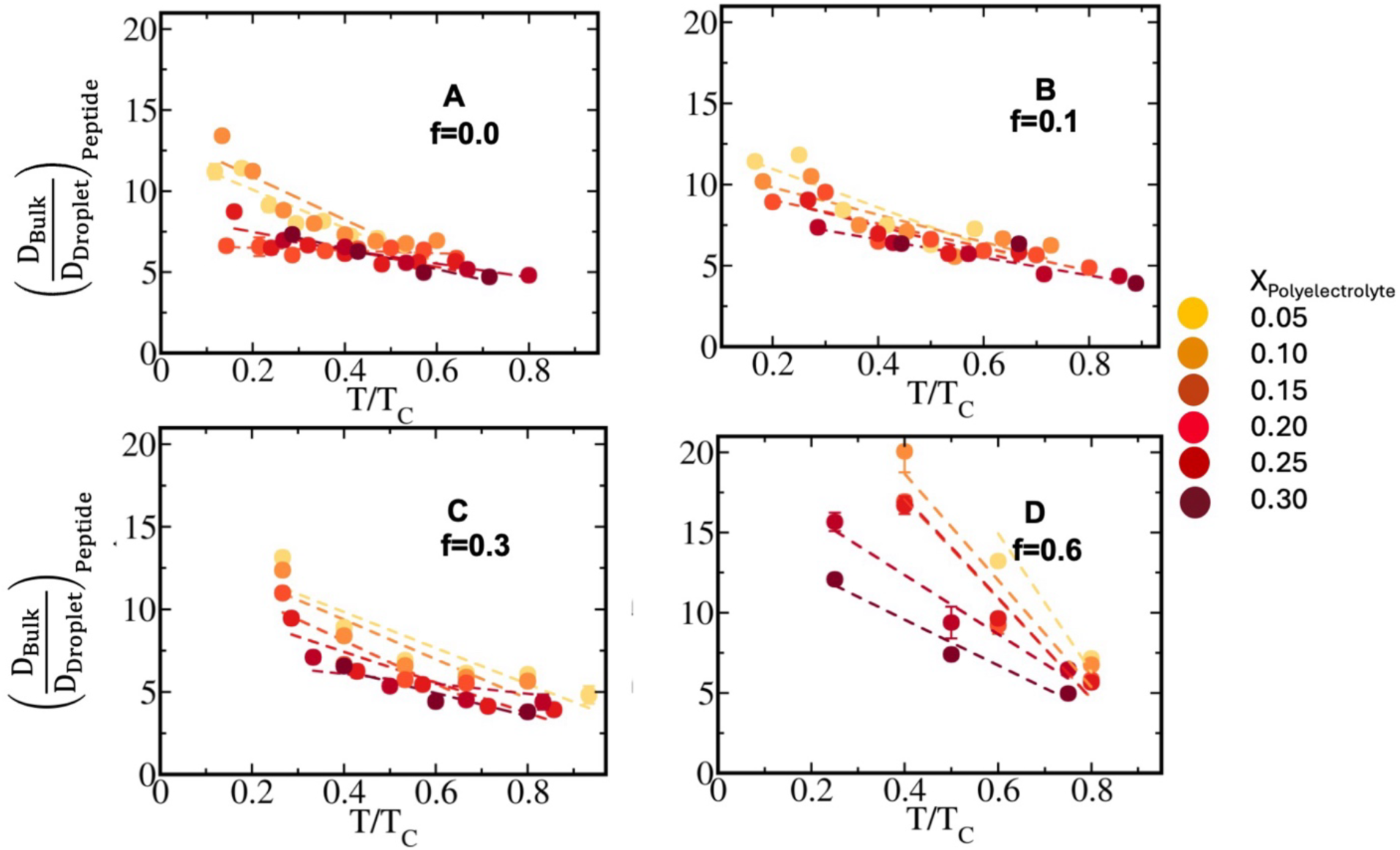
Differential dynamics of peptides in condensate phase and bulk. Ratio of average diffusivity of peptides in condensate phase and bulk 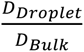 along temperature scaled to criticality for sequence pairs (A) f=0.0 and (U)_20_, (B) f=0.1 and (U)_20_ and (C) f=0.3 and (U)_20_ (D) f=0.6 and (U)_20_. Different mixing fractions has been shown in each panel ranging from *X_polyelectrolyte_* = 0.05 − 0.30 with yellow to dark red.

In addition, to connect translational dynamics to chain’s internal motions, we have computed peptide’s end-to-end distance dynamics to elucidate chain reconfiguration timescales as follows:

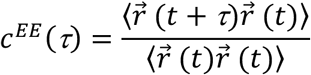

where 𝑟⃗ (𝑡) represents end-to-end distance vector of peptides (Fig 8 and convergence of correlation functions has been shown in Fig S8 at T/Tc=0.6 for all the designed peptide condensates at X_Polyelectrolyte_=0.15 from independent trajectories).

**Fig 8.**
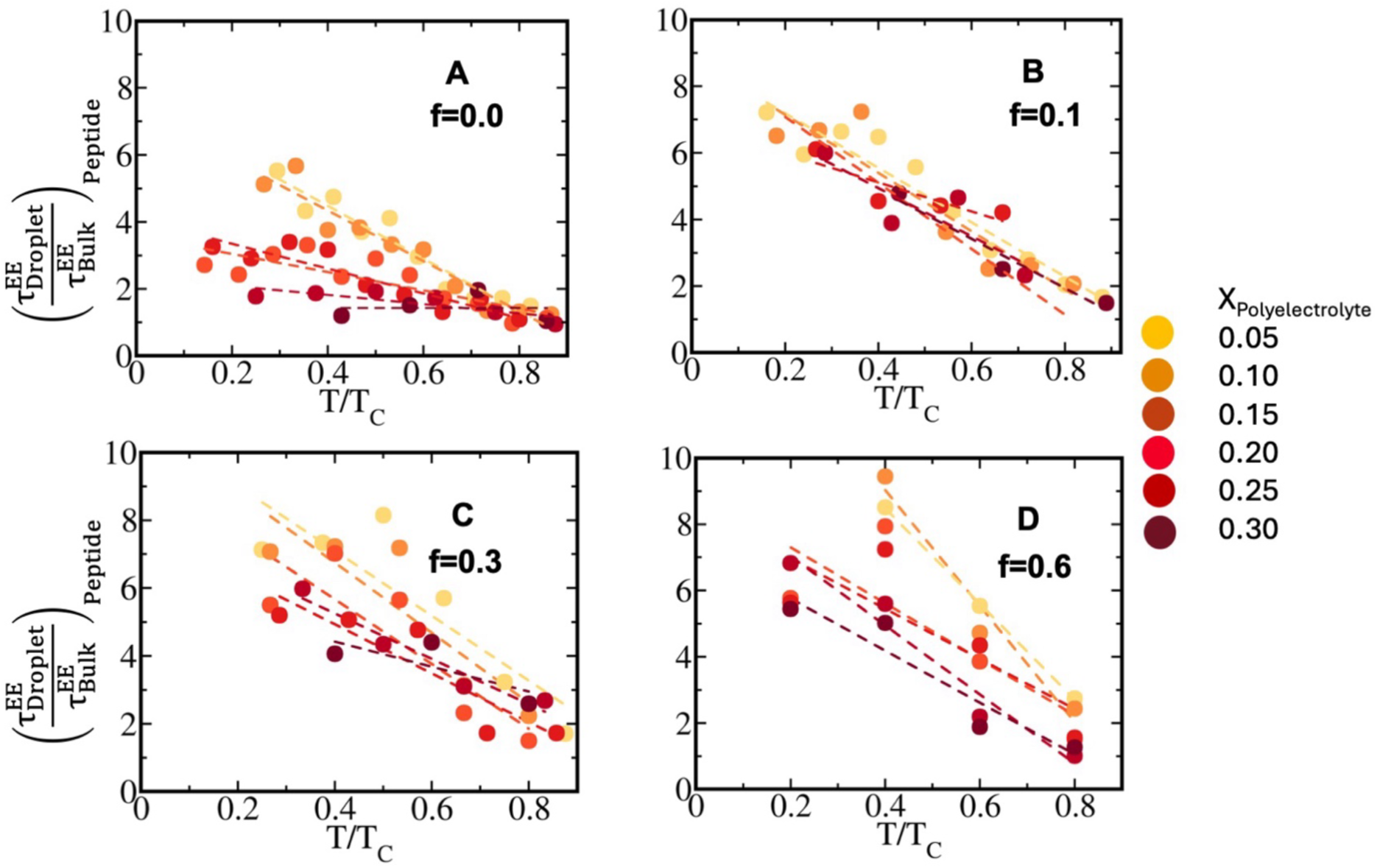
Peptide chain reconfiguration dynamics in condensates. Average Chain reconfiguration timescale of peptides compared to bulk 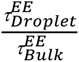 along temperature scaled to critical temperature for droplets of sequence pairs (A) f=0.0 and (U)_20_, (B) f=0.1 and (U)_20_ and (C) f=0.3 and (U)_20_ (D) f=0.6 and (U)_20_. Different mixing fractions has been shown in each panel ranging from *X_polyelectrolyte_* = 0.05 − 0.30 with yellow to dark red. Convergence of correlation functions has been shown in Fig S8 at T/Tc=0.6 for all the designed peptide condensates at X_Polyelectrolyte_=0.15 from independent trajectories

For purely electrostatic sequences (𝑓 = 0.0), peptide diffusivity in bulk is nearly 15-times higher than in the droplet phase at low polyelectrolyte mixing (X_Polyelectrolyte_=0.05) and far from criticality (T/T_C_=0.3). Increasing polyelectrolyte enrichment enhances droplet diffusivity (𝐷_Droplet_; Fig. 7A) by significantly weakening dense-phase interactions (Figs. 2A, 3A), reducing the bulk–droplet contrast to ∼5-times. Until X_Polyelectrolyte_=0.10 as the droplet retains its stability (as observed in Fig. 4A and B), the dense network showcase slower droplet dynamics relative to bulk (i.e: 12-15 times slower than bulk) far away from criticality (T/Tc=0.3). An abrupt drop in the ratio (i.e: 6-8 times slower than bulk) has been observed at same distance from criticality once X_Polyelectrolyte_ is beyond 0.15 correlated to depleted stability. As the temperature approaches the critical point, 𝐷_Droplet_ increases monotonically, with droplet diffusivity remaining ∼5-times slower than bulk near criticality.

Clear signature of polyelectrolyte mixing fraction mediated enhanced diffusion of peptides is evident in electrostatic dominated condensates than hydrophobic ones. This may originate from long-range and exchangeable electrostatic attractions buffer strong polyelectrolyte repulsions while the same is sticky in hydrophobic condensates having primarily short-range attractions. Consistent with translational diffusion, end-to-end distance vector dynamics of peptides in the droplet phase are ∼6 times slower than in bulk at low polyelectrolyte fraction (X_Polyelectrolyte_=0.05) and far from criticality (T/T_C_=0.3), but this disparity diminishes with increasing polyelectrolyte content. Upon approaching the critical regime, the ratio 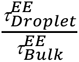 decreases monotonically and approaches unity, indicating comparable chain conformational dynamics in the droplet and bulk phases at single molecule level (Fig. 8A).

At low sequence hydrophobicity (f=0.1), we observe no measurable effect of polyelectrolyte fraction on droplet–bulk diffusivity ratio. Far from criticality (T/T_C_=0.2), droplet diffusivity is ∼8–12 times slower than bulk across all polyelectrolyte mixing fractions. With increasing temperature, the ratio 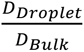 decreases monotonically, reaching ∼5 times slowdown near the critical regime. Similarly, end-to-end vector dynamics is also insensitive to polyelectrolyte enrichment: the reconfiguration timescale in droplets is ∼8 times that of bulk at T/T_C_=0.3 , decreasing monotonically to ∼1.5-times as the temperature approaches the critical point (Fig. 8B).

At moderate sequence hydrophobicity (f=0.3; Fig 7C), peptides again exhibit pronounced translational slowdown at low polyelectrolyte fraction (X_Polyelectrolyte_=0.05). For f=0.3, the diffusivity ratio 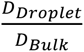 prominently showcase effect of polyelectrolyte content. At T/T_C_=0.3 , bulk polymers diffuse ∼14-times faster than in the dense phase. In contrast, at higher polyelectrolyte enrichment and elevated temperatures, droplet peptides are only ∼5-times slower than bulk, indicating increasingly liquid-like behaviour (Fig. 7C). Consistently, end-to-end vector reconfiguration timescales in the droplet phase are ∼4–8 times longer than in bulk, depending on polyelectrolyte mole fraction, showcasing enhanced slowdown of reconfigurational dynamics at higher hydrophobicity and decrease monotonically to ∼1.5–2 times as the temperature approaches the critical point (Fig. 8C).

For highly hydrophobic sequences (f=0.6), bulk diffusivity is ∼15–20 times higher than droplet diffusivity at low polyelectrolyte fractions (X_Polyelectrolyte_=0.05-0.15) and far from criticality (T/T_C_=0.3), indicating a solid-like assembly dominated by short-range hydrophobic interactions. At higher polyelectrolyte enrichment (X_Polyelectrolyte_=0.30), this contrast reduces to ∼10-times signifying fairly less liquid-like character. With increasing temperature, the bulk–droplet diffusivity ratio decreases monotonically, reaching ∼5 times near the critical regime (Fig 7D). Consistent with translational diffusion, end-to-end vector dynamics in the dense phase are ∼10 times slower than in bulk at low polyelectrolyte fraction and T/Tc =0.3, and ∼6× slower at higher polyelectrolyte concentrations (Fig 8D). 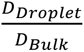 has been shown at constant T*=0.3 for all the sequence variants along polyelectrolyte composition (Fig S9). Mean squared displacements (Fig S10) and end-to-end distance time correlation function of peptides (Fig S11) in condensate phase been shown as a function of polyelectrolyte composition for an absolute T*=0.3.

### Droplet Shape Anisotropy and correlated sticker contact lifetimes

To correlate the stability pattern of condensate phase to the geometry of the same, we estimate deviation of the shape of condensates from a sphere encompassing the same. Droplet shape anisotropy (p) has been defined as:

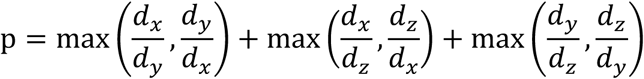

where 𝑑_𝑥_, 𝑑_𝑦_, 𝑑_𝑧_ are the largest diameter in each dimension of the largest cluster. An ideal sphere should have a value of 3 along this definition and deviation from the same indicates a distorted condensate. Phase separated condensates tend to acquire a spherical shape to reduce surface tension. Deviation of shape of a condensate from a spherical one may originate from competitive interactions. Fig 9 plots the anisotropy of condensate phase for designed sequences with varying hydrophobicity as a function of polyelectrolyte mixing fraction (Fig 9 panel A) at constant T/Tc=0.4 and along temperature scaled to criticality at constant polyelectrolyte mixing fraction X_Polyelectrolyte_=0.20 (Fig 9 panel B). With increase in polyelectrolyte’s concentration, droplet shape anisotropy increases gradually. However, the slope decreases with enhancement of sequence hydrophobicity consistent with stability analysis in Fig 4A and B showcasing a dense near spherical network scaffolded by polyelectrolytes. Sequence having dominant electrostatic interactions form condensates showcase shape anisotropy 3.6 at X_Polyelectrolyte_=0.05, the same enhances to 4.5 at X_Polyelectrolyte_=0.30 indicating weaking of condensate sticker interactions with fusion of polyelectrolytes as observed in Fig 2A and Fig 3A. Extremely hydrophobic sequences form condensates with nearly uniform shape anisotropy 3.7 -3.9 at various polyelectrolyte mixing fractions denoting a stabilizing effect of RNA-like polyelectrolytes on droplet phase due to availability of a hub of short-range interaction sites on such polyelectrolytes promoted by hydrophobicity of the peptide sequence. The condensates are near spherical due to tight short-range peptide-peptide and peptide-polyelectrolyte interactions. Along temperature scaled to criticality, droplet shape anisotropy enhances as the stability of the dense phase diminish for dominant electrostatic interaction driven condensates within the temperature window of T/Tc =0.2 to 0.7 while the droplet retains near spherical shape even at enhanced temperatures for hydrophobic sequences (Fig. 9B and Fig. S12).

**Fig 9.**
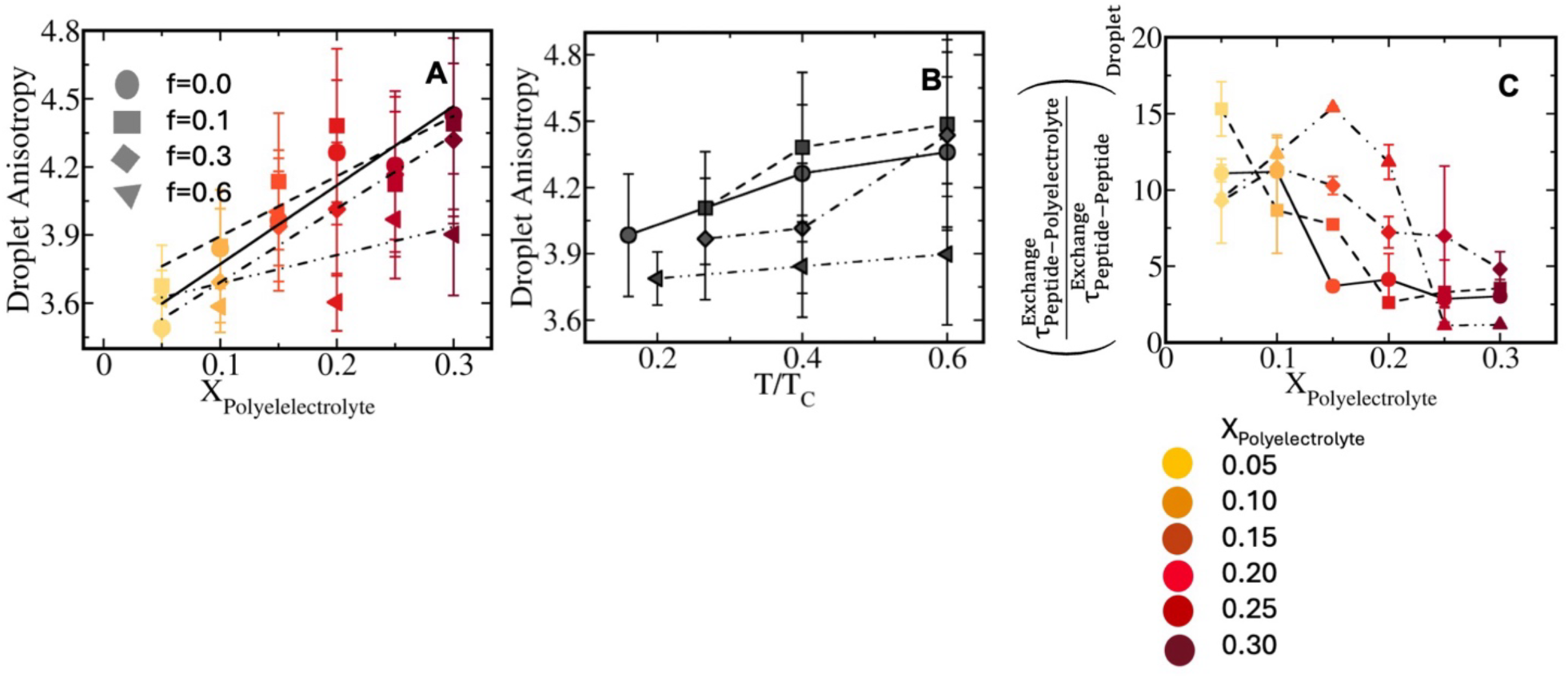
Anisotropy in geometry of condensate. **(A)** Shape anisotropy parameter shown along *X_polyelectrolyte_* at fixed T/Tc=0.4. (B) Shape anisotropy parameter along scaled temperature to criticality in the range 0.2-0.7 for condensates formed by sequence pairs. f=0.0 and (U)_20_ (shown with circle, connected with solid line), f=0.1 and (U)_20_ (shown with squares, connected with dashed line), f=0.3 and (U)_20_ (shown with diamond shapes, connected with dot dashed line) f=0.6 and (U)_20_ (shown with triangles, connected with double dot dashed line) at *X_polyelectrolyte_* = 0.15. Color codes represent polyelectrolyte mixing fractions. (C) Ratio of average lifetimes of peptide-polyelectrolyte contact lifetimes with respect peptide-peptide pairs 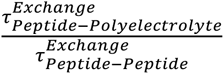 along polyelectrolyte mixing fraction for sequence pairs f=0.0 and (U)_20_ (shown with circle connected with solid line), f=0.1 and (U)_20_ (shown with squares, connected with dashed line), f=0.3 and (U)_20_ (shown with diamond shapes, connected with dot dashed line) f=0.6 and (U)_20_ (shown with triangles, connected with double dot dashed line).

To correlate stability and shape of the condensates to a microscopic understanding of sticker exchange dynamics in network, we have introduced a sticker exchange time correlation function among peptides, peptide-polyelectrolyte and polyelectrolytes pairs in dense phase as

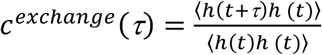

h(t) denotes a step function; h(t)=1 when two polymers (either peptide or polyelectrolytes) are neighboring ones for a time frame and 0 otherwise. Time correlation functions has been plotted for peptide-peptide (P-P), peptide-polyelectrolyte (P-PE) and polyelectrolyte-polyelectrolyte (PE-PE) neighbors in Fig S13.

Ratio of average lifetimes of peptide-polyelectrolyte contact lifetimes with respect to peptide-peptide pairs has been shown in Fig 9C for various designed sequences along polyelectrolyte mixing fraction at a constant distance from criticality (T/Tc=0.6).

Peptide-polyelectrolyte interactions are nearly 10-15 times longer than the same among peptide-peptide pairs at low X_Polyelectrolyte_ and decreases with increased polyelectrolyte mixing fraction. Beyond X_Polyelectrolyte_=0.15, 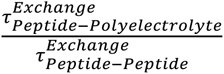 showcase gradual depletion and decays fairly rapidly at near extreme polyelectrolyte mixing fractions. Such observation indicates long-lived peptide-polyelectrolyte interactions act as the scaffold for droplet formation while easily exchangeable peptide-peptide ones stabilize largescale peptide assembly. For an extremely hydrophobic sequence, 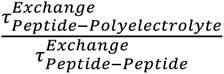 showcase a maximum around X_Polyelectrolyte_=0.10-0.15 indicating an intermediate polyelectrolyte concentration maintains significant contact lifetime among peptide and polyelectrolyte that in turn stabilizes droplet with exchangeable peptide-peptide interactions.

Visual representation of structural morphology and droplet shape has been shown in Fig. 10 for two extreme polyelectrolytes mixing fractions X_Polyelectrolyte_ =0.05 and 0.25 and for dominant electrostatic (Fig. 10 A and B) and hydrophobic condensates at the same distance from criticality (Fig. 10 C-D). It is very interesting to note that at dominant electrostatic regime of sequences, peptides and polyelectrolytes form a dense network at lower polyelectrolyte limit (X_Polyelectrolyte_ =0.05), while smaller droplets are connected by polyelectrolyte-peptide cross links at relatively higher polyelectrolyte concentration leading weaker stability of the mesh. Droplets formed by dominant hydrophobic interactions are dense and spherical irrespective of polyelectrolyte concentration and showcase polyelectrolytes acting as hubs of short-range sites while polyelectrolyte-polyelectrolyte repulsion has been buffered by intermittent peptides. Differential diffusivity obtained in Fig. 6 and end-to-end distance vector dynamics of peptides in Fig 8 are essentially connected to such structural morphology and stability of condensates.

**Fig 10.**
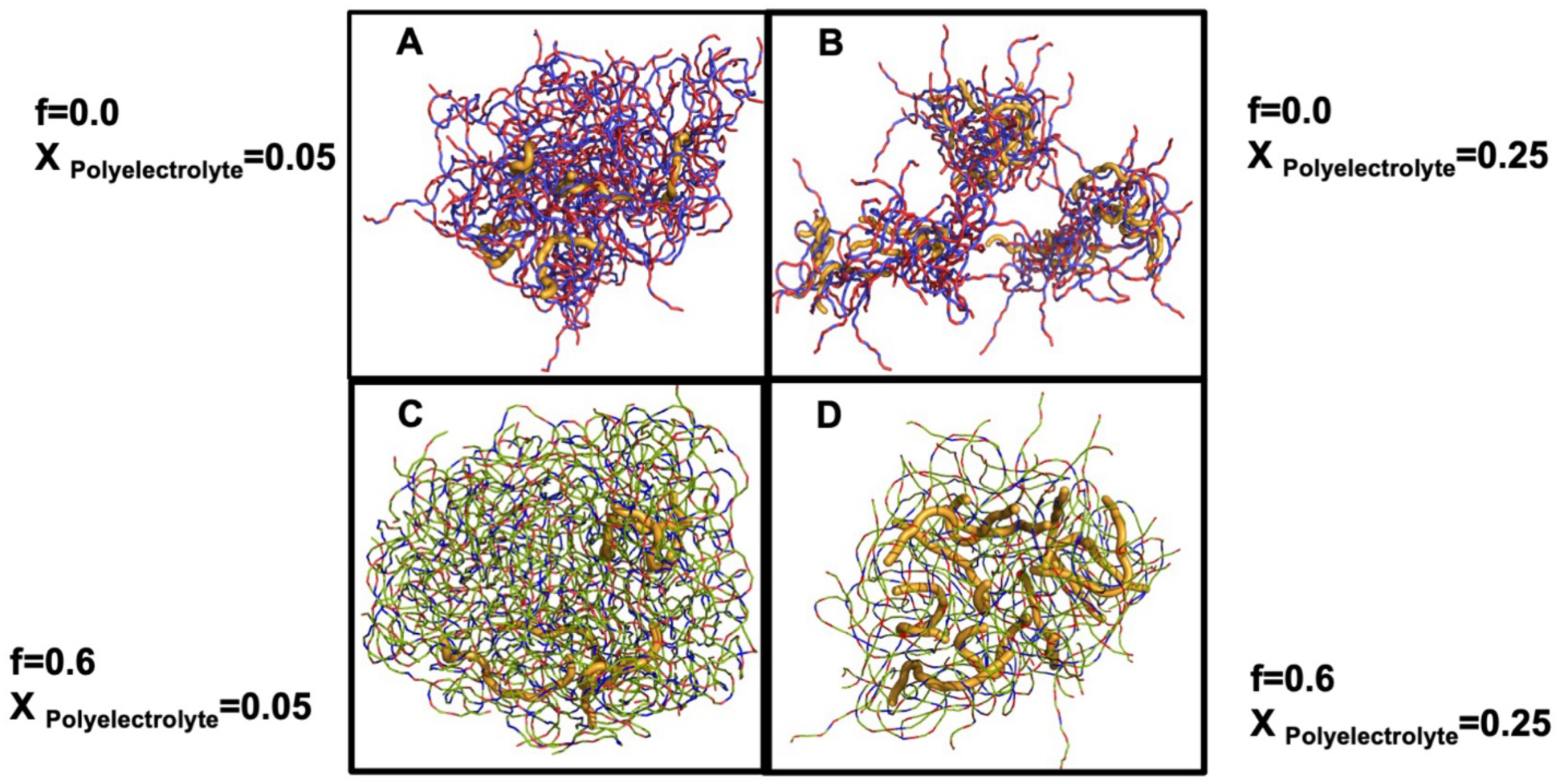
Representative snapshots of condensates delineating structural morphology and related geometric features for (A) lower population of polyelectrolytes *X_polyelectrolyte_* = 0.05 , (B) higher population of polyelectrolytes *X_polyelectrolyte_* = 0.25 for electrostatics dominated condensate with sequence pair f=0.0 and (U)_20_ (C) lower population of polyelectrolytes *X_polyelectrolyte_* = 0.05 (D) higher population of polyelectrolytes *X_polyelectrolyte_* = 0.25 for hydrophobic interaction dominated condensates with sequence pair f=0.6 and (U)_20_. Positively charged amino acid residues has been shown with blue, negative ones with red and neutral hydrophobic ones has been shown with limon green. Polyelectrolytes has been colored with orange.

## Conclusion

Heterotypic condensates play an important role in spatio-temporal organization of cellular material for tuning minute biological regulations. Often such condensates are formed by proteins and RNAs at various stoichiometries. RNA being the hub of both long and short-range interactions intricately determine the liquid-like nature of droplets and stability. To elucidate a molecular level picture of the same, present study explores RNA-like polyelectrolyte’s influence in altering stability and dynamics of heterotypic condensates formed by a series of sequences having a spectrum of long-to-short range interactions and having various polyelectrolyte compositions. A systematic enhancement of short-range hydrophobic interactions in lieu of long-range electrostatics in designed sequences showcase a dual role of such polyelectrolytes in stabilizing or destabilizing condensates. Excessive polyelectrolyte concentration depletes condensate’s stability rapidly at dominant electrostatic limit while such condensates are stable at intermediate polyelectrolyte concentration. Polyelectrolytes on the other hand acts as a stabilizing sticker hub for condensates dominated by extreme hydrophobic sequences. Translational diffusivity of RNA like polyelectrolytes in condensate is nearly 20 to 40% higher than polyelectrolytes in dominant electrostatic limit whereas polyelectrolyte diffusion is nearly 50% slower than peptides in extremely hydrophobic condensates pointing to a scaffold like behaviour. Peptide’s diffusivity in droplet phase compared to bulk ranges from 20-times to 5-times respectively upon enhancement of polyelectrolyte composition subject to various designed sequence hydrophobicities. Enrichment of polyelectrolytes in condensate phase saturates at largest mole fraction 0.2 to 0.25 and showcase non-linearity. Polyelectrolytes being the hub of long and short-range interaction sites in hydrophobic condensates act as stickers and peptide-polyelectrolyte interactions form the core of droplet stability whereas peptide-peptide interactions enhance the peptide enrichment in droplets. Long-range repulsions diminish stability in condensates dominated by electrostatic interactions at excessive polyelectrolyte concentration. Droplet shape tends toward spherical ones at lower polyelectrolyte limit having marked deviation from spherical shape for dominant electrostatics condensates at excessive polyelectrolyte concentration. Overall, the study depicts opposing tuneable roles of RNA-like polyelectrolytes on stability-dynamics paradigm of condensates having a spectrum of long-to short range interactions. Present study creates a standpoint upon which we aim to quantify the precise effect of cation-π cross interactions that will compete with both electrostatic and hydrophobic interactions and will tune stability-dynamics paradigm of protein-polyelectrolyte condensates. Our upcoming study aims to uncover this competitive role of cation-π interactions over presently described dwell of electrostatic and hydrophobic interactions modulated by polyelectrolytes completing the picture of real protein-RNA condensates. Although present study incorporates only dominant electrostatic and hydrophobic interactions and does not have the effect of cation-π interactions, the outcomes will be crucial to quantitatively measure the precise effect of cation-π interactions on condensate’s stability and dynamics being the reference hereafter.

## Supporting Information

Dissection of interaction energy into self and cross components in peptide–polyelectrolyte condensates for sequences designed with various hydrophobicity (Fig S1-4), convergence of diffusivity of peptides (Fig S5) and polyelectrolytes (Fig S6) in condensate phase from different independent simulation runs, Comparative mean squared displacement (MSD) of peptides in droplet and bulk phases at fixed distance from criticality for various sequences hydrophobicity at intermediate polyelectrolyte composition (Fig S7), convergence of end-to-end distance vector time correlation for peptides from different independent simulation runs (Fig S8) at fixed distance from criticality and intermediate polyelectrolyte composition, 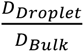 has been shown at constant T*=0.3 for all the sequence variants along polyelectrolyte composition (Fig S9). Mean squared displacements (Fig S10) and end-to-end distance time correlation function of peptides (Fig S11) in condensate phase been shown as a function of polyelectrolyte composition for an absolute T*=0.3, Droplet shape anisotropy along temperature and polyelectrolyte composition for various designed sequence hydrophobicity (Fig S12), Neighbor exchange time correlation functions has been plotted for peptide-peptide (P-P), peptide-polyelectrolyte (P-PE) and polyelectrolyte-polyelectrolyte (PE-PE) pairs at various sequence hydrophobicity at a fixed distance from criticality in Fig S13.

## References

(1) Qiao, Y.; Zia, A.; Wu, G.; Liu, Z.; Guo, J.; Chu, M.; He, H.; Wang, F.; Xu, B. Context-Dependent Heterotypic Assemblies of Intrinsically Disordered Peptides. J. Am. Chem. Soc. 2025. 10.1021/jacs.4c12150.

(2) Agarwal, A.; Arora, L.; Rai, S. K.; Avni, A.; Mukhopadhyay, S. Spatiotemporal Modulations in Heterotypic Condensates of Prion and α-Synuclein Control Phase Transitions and Amyloid Conversion. Nat. Commun. 2022, 13 (1). 10.1038/s41467-022-28797-5.

(3) Rai, S. K.; Khanna, R.; Avni, A.; Mukhopadhyay, S. Heterotypic Electrostatic Interactions Control Complex Phase Separation of Tau and Prion into Multiphasic Condensates and Co-Aggregates. Proc. Natl. Acad. Sci. U. S. A. 2023, 120 (2). 10.1073/pnas.2216338120.

(4) Hazra, M. K. Differential Sequence Charge Clustering and Mixing Ratio Affect Stability and Dynamics of Heterotypic Peptide Condensates. Physical Chemistry Chemical Physics 2026, 28 (1), 692–706. 10.1039/d5cp03436a.

(5) Musacchio, A. On the Role of Phase Separation in the Biogenesis of Membraneless Compartments. EMBO J. 2022, 41 (5). 10.15252/embj.2021109952.

(6) Kedersha, N.; Ivanov, P.; Anderson, P. Stress Granules and Cell Signaling: More than Just a Passing Phase? Trends in Biochemical Sciences. October 2013, pp 494–506. 10.1016/j.tibs.2013.07.004.

(7) Gao, G.; Sumrall, E. R.; Walter, N. G. Nanoscale Domains Govern Local Diffusion and Ageing within Fused-in-Sarcoma Condensates. Nat. Nanotechnol. 2025. 10.1038/s41565-025-02077-x.

(8) Wadsworth, G.; Srinivasan, S.; Lai, L.; Datta, M.; Gopalan, V.; Banerjee, P. R. RNA-Driven Phase Transitions in Biomolecular Condensates. Molecular Cell. Cell Press October 3, 2024, pp 3692–3705. 10.1016/j.molcel.2024.09.005.

(9) Roden, C.; Gladfelter, A. S. RNA Contributions to the Form and Function of Biomolecular Condensates. Nature Reviews Molecular Cell Biology. Nature Research March 1, 2021, pp 183–195. 10.1038/s41580-020-0264-6.

(10) Ladouceur, A.-M.; Singh Parmar, B.; Biedzinski, S.; Wall, J.; Graydon Tope, S.; Cohn, D.; Kim, A.; Soubry, N.; Reyes-Lamothe, R.; Weber, S. C.; Young, R. A. Clusters of Bacterial RNA Polymerase Are Biomolecular Condensates That Assemble through Liquid-Liquid Phase Separation. 10.1073/pnas.2005019117/-/DCSupplemental.

(11) Monterroso, B.; Zorrilla, S.; Sobrinos-Sanguino, M.; Robles-Ramos, M. A.; López-Álvarez, M.; Margolin, W.; Keating, C. D.; Rivas, G. Bacterial FtsZ Protein Forms Phase-separated Condensates with Its Nucleoid-associated Inhibitor SlmA. EMBO Rep. 2019, 20 (1). 10.15252/embr.201845946.

(12) Cubuk, J.; Alston, J. J.; Incicco, J. J.; Singh, S.; Stuchell-Brereton, M. D.; Ward, M. D.; Zimmerman, M. I.; Vithani, N.; Griffith, D.; Wagoner, J. A.; Bowman, G. R.; Hall, K. B.; Soranno, A.; Holehouse, A. S. The SARS-CoV-2 Nucleocapsid Protein Is Dynamic, Disordered, and Phase Separates with RNA. Nat. Commun. 2021, 12 (1). 10.1038/s41467-021-21953-3.

(13) Jing, H.; Korasick, D. A.; Emenecker, R. J.; Morffy, N.; Wilkinson, E. G.; Powers, S. K.; Strader, L. C. Regulation of AUXIN RESPONSE FACTOR Condensation and Nucleo-Cytoplasmic Partitioning. Nat. Commun. 2022, 13 (1). 10.1038/s41467-022-31628-2.

(14) Alshareedah, I.; Moosa, M. M.; Raju, M.; Potoyan, D. A.; Banerjee, P. R. Phase Transition of RNA−protein Complexes into Ordered Hollow Condensates. Proc. Natl. Acad. Sci. U. S. A. 2020, 117 (27), 15650–15658. 10.1073/pnas.1922365117.

(15) Molliex, A.; Temirov, J.; Lee, J.; Coughlin, M.; Kanagaraj, A. P.; Kim, H. J.; Mittag, T.; Taylor, J. P. Phase Separation by Low Complexity Domains Promotes Stress Granule Assembly and Drives Pathological Fibrillization. Cell 2015, 163 (1), 123–133. 10.1016/j.cell.2015.09.015.

(16) Patel, A.; Lee, H. O.; Jawerth, L.; Maharana, S.; Jahnel, M.; Hein, M. Y.; Stoynov, S.; Mahamid, J.; Saha, S.; Franzmann, T. M.; Pozniakovski, A.; Poser, I.; Maghelli, N.; Royer, L. A.; Weigert, M.; Myers, E. W.; Grill, S.; Drechsel, D.; Hyman, A. A.; Alberti, S. A Liquid-to-Solid Phase Transition of the ALS Protein FUS Accelerated by Disease Mutation. Cell 2015, 162 (5), 1066–1077. 10.1016/j.cell.2015.07.047.

(17) Alshareedah, I.; Borcherds, W. M.; Cohen, S. R.; Singh, A.; Posey, A. E.; Farag, M.; Bremer, A.; Strout, G. W.; Tomares, D. T.; Pappu, R. V.; Mittag, T.; Banerjee, P. R. Sequence-Specific Interactions Determine Viscoelasticity and Ageing Dynamics of Protein Condensates. Nat. Phys. 2024, 20 (9), 1482–1491. 10.1038/s41567-024-02558-1.

(18) Mittag, T.; Pappu, R. V. A Conceptual Framework for Understanding Phase Separation and Addressing Open Questions and Challenges. Molecular Cell. Cell Press June 16, 2022, pp 2201–2214. 10.1016/j.molcel.2022.05.018.

(19) Linsenmeier, M.; Faltova, L.; Morelli, C.; Capasso Palmiero, U.; Seiffert, C.; Küffner, A. M.; Pinotsi, D.; Zhou, J.; Mezzenga, R.; Arosio, P. The Interface of Condensates of the HnRNPA1 Low-Complexity Domain Promotes Formation of Amyloid Fibrils. Nat. Chem. 2023, 15 (10), 1340–1349. 10.1038/s41557-023-01289-9.

(20) Jawerth, L.; Fischer-Friedrich, E.; Saha, S.; Wang, J.; Franzmann, T.; Zhang, X.; Sachweh, J.; Ruer, M.; Ijavi, M.; Saha, S.; Mahamid, J.; Hyman, A. A.; Jülicher, F. Protein Condensates as Aging Maxwell Fluids. https://www.science.org.

(21) Alshareedah, I.; Kaur, T.; Ngo, J.; Seppala, H.; Kounatse, L. A. D.; Wang, W.; Moosa, M. M.; Banerjee, P. R. Interplay between Short-Range Attraction and Long-Range Repulsion Controls Reentrant Liquid Condensation of Ribonucleoprotein-RNA Complexes. J. Am. Chem. Soc. 2019, 141 (37), 14593–14602. 10.1021/jacs.9b03689.

(22) Banerjee, P. R.; Milin, A. N.; Moosa, M. M.; Onuchic, P. L.; Deniz, A. A. Reentrant Phase Transition Drives Dynamic Substructure Formation in Ribonucleoprotein Droplets. Angewandte Chemie - International Edition 2017, 56 (38), 11354–11359. 10.1002/anie.201703191.

(23) Laghmach, R.; Alshareedah, I.; Pham, M.; Raju, M.; Banerjee, P. R.; Potoyan, D. A. RNA Chain Length and Stoichiometry Govern Surface Tension and Stability of Protein-RNA Condensates. iScience 2022, 25 (4). 10.1016/j.isci.2022.104105.

(24) Fei, J.; Jadaliha, M.; Harmon, T. S.; Li, I. T. S.; Hua, B.; Hao, Q.; Holehouse, A. S.; Reyer, M.; Sun, Q.; Freier, S. M.; Pappu, R. V.; Prasanth, K. V.; Ha, T. Quantitative Analysis of Multilayer Organization of Proteins and RNA in Nuclear Speckles at Super Resolution. J. Cell Sci. 2017, 130 (24), 4180–4192. 10.1242/jcs.206854.

(25) Yamazaki, T.; Souquere, S.; Chujo, T.; Kobelke, S.; Chong, Y. S.; Fox, A. H.; Bond, C. S.; Nakagawa, S.; Pierron, G.; Hirose, T. Functional Domains of NEAT1 Architectural LncRNA Induce Paraspeckle Assembly through Phase Separation. Mol. Cell 2018, 70 (6), 1038–1053.e7. 10.1016/j.molcel.2018.05.019.

(26) Takakuwa, H.; Yamazaki, T.; Souquere, S.; Adachi, S.; Yoshino, H.; Fujiwara, N.; Yamamoto, T.; Natsume, T.; Nakagawa, S.; Pierron, G.; Hirose, T. Shell Protein Composition Specified by the LncRNA NEAT1 Domains Dictates the Formation of Paraspeckles as Distinct Membraneless Organelles. Nat. Cell Biol. 2023, 25 (11), 1664–1675. 10.1038/s41556-023-01254-1.

(27) Quinodoz, S. A.; Guttman, M. Essential Roles for RNA in Shaping Nuclear Organization. Cold Spring Harb. Perspect. Biol. 2022, 14 (5). 10.1101/cshperspect.a039719.

(28) Arun, G.; Aggarwal, D.; Spector, D. L. MALAT1 Long Non-Coding RNA: Functional Implications. Non-coding RNA. MDPI AG June 1, 2020. 10.3390/NCRNA6020022.

(29) Shevtsov, S. P.; Dundr, M. Nucleation of Nuclear Bodies by RNA. Nat. Cell Biol. 2011, 13 (2), 167–173. 10.1038/ncb2157.

(30) Chen, X.; Fansler, M. M.; Janjoš, U.; Ule, J.; Mayr, C. The FXR1 Network Acts as a Signaling Scaffold for Actomyosin Remodeling. Cell 2024, 187 (18), 5048–5063.e25. 10.1016/j.cell.2024.07.015.

(31) Sanchez-Burgos, I.; Espinosa, J. R.; Joseph, J. A.; Collepardo-Guevara, R. RNA Length Has a Non-Trivial Effect in the Stability of Biomolecular Condensates Formed by RNA-Binding Proteins. PLoS Comput. Biol. 2022, 18 (2). 10.1371/journal.pcbi.1009810.

(32) Edelstein, I.; Levy, Y. How the Extent of Protein Folding and Oligomerization Modulate Condensate Formation and Properties. Journal of Physical Chemistry Letters 2025, 16, 11248–11258. 10.1021/acs.jpclett.5c02083.

(33) Tejedor, A. R.; Sanchez-Burgos, I.; Estevez-Espinosa, M.; Garaizar, A.; Collepardo-Guevara, R.; Ramirez, J.; Espinosa, J. R. Protein Structural Transitions Critically Transform the Network Connectivity and Viscoelasticity of RNA-Binding Protein Condensates but RNA Can Prevent It. Nat. Commun. 2022, 13 (1). 10.1038/s41467-022-32874-0.

(34) Garcia-Jove Navarro, M.; Kashida, S.; Chouaib, R.; Souquere, S.; Pierron, G.; Weil, D.; Gueroui, Z. RNA Is a Critical Element for the Sizing and the Composition of Phase-Separated RNA–Protein Condensates. Nat. Commun. 2019, 10 (1), 1–13. 10.1038/s41467-019-11241-6.

(35) Yamamoto, T.; Yamazaki, T.; Ninomiya, K.; Hirose, T. Nascent Ribosomal RNA Act as Surfactant That Suppresses Growth of Fibrillar Centers in Nucleolus. *Commun*. Biol. 2023, 6 (1). 10.1038/s42003-023-05519-1.

(36) Alshareedah, I.; Muhammad Moosa, M.; Raju, M.; Potoyan, D. A.; Banerjee, P. R. Phase Transition of RNA−protein Complexes into Ordered Hollow Condensates. 2020, 117, 15650–15658. 10.1073/pnas.1922365117/-/DCSupplemental.

(37) Alshareedah, I.; Moosa, M. M.; Banerjee, P. R. Programmable Viscoelasticity in Protein-RNA Condensates with Disordered Sticker-Spacer Polypeptides. bioRxiv 2021, 2021.01.24.427968.

(38) Alshareedah, I.; Thurston, G. M.; Banerjee, P. R. Quantifying Viscosity and Surface Tension of Multicomponent Protein-Nucleic Acid Condensates. Biophys. J. 2021, 120 (7), 1161–1169. 10.1016/j.bpj.2021.01.005.

(39) Alshareedah, I.; Moosa, M. M.; Pham, M.; Potoyan, D. A.; Banerjee, P. R. Programmable Viscoelasticity in Protein-RNA Condensates with Disordered Sticker-Spacer Polypeptides. Nat. Commun. 2021, 12 (1). 10.1038/s41467-021-26733-7.

(40) Alshareedah, I.; Moosa, M. M.; Pham, M.; Potoyan, D. A.; Banerjee, P. R. Programmable Viscoelasticity in Protein-RNA Condensates with Disordered Sticker-Spacer Polypeptides. Nat. Commun. 2021, 12 (1). 10.1038/s41467-021-26733-7.

(41) Alshareedah, I.; Muhammad Moosa, M.; Raju, M.; Potoyan, D. A.; Banerjee, P. R. Phase Transition of RNA−protein Complexes into Ordered Hollow Condensates. 2020, 117, 15650–15658. 10.1073/pnas.1922365117/-/DCSupplemental.

(42) Alshareedah, I.; Moosa, M. M.; Pham, M.; Potoyan, D. A.; Banerjee, P. R. Programmable Viscoelasticity in Protein-RNA Condensates with Disordered Sticker-Spacer Polypeptides. Nat. Commun. 2021, 12 (1). 10.1038/s41467-021-26733-7.

(43) Maharana, S.; Wang, J.; Papadopoulos, D. K.; Richter, D.; Pozniakovsky, A.; Poser, I.; Bickle, M.; Rizk, S.; Guillén-Boixet, J.; Franzmann, T. M.; Jahnel, M.; Marrone, L.; Chang, Y.-T.; Sterneckert, J.; Tomancak, P.; Hyman, A. A.; Alberti, † Simon. MOLECULAR BIOLOGY RNA Buffers the Phase Separation Behavior of Prion-like RNA Binding Proteins. https://www.science.org.

(44) Aarum, J.; Cabrera, C. P.; Jones, T. A.; Rajendran, S.; Adiutori, R.; Giovannoni, G.; Barnes, M. R.; Malaspina, A.; Sheer, D. Enzymatic Degradation of RNA Causes Widespread Protein Aggregation in Cell and Tissue Lysates . EMBO Rep. 2020, 21 (10). 10.15252/embr.201949585.

(45) Mathieu, C.; Pappu, R. V; Taylor, J. P. Beyond Aggregation: Pathological Phase Transitions in Neurodegenerative Disease. https://www.science.org.

(46) Bokros, M.; Balukoff, N. C.; Grunfeld, A.; Sebastiao, M.; Beurel, E.; Bourgault, S.; Lee, S. RNA Tailing Machinery Drives Amyloidogenic Phase Transition. Proc. Natl. Acad. Sci. U. S. A. 2024, 121 (23). 10.1073/pnas.2316734121.

(47) Morelli, C.; Faltova, L.; Capasso Palmiero, U.; Makasewicz, K.; Papp, M.; Jacquat, R. P. B.; Pinotsi, D.; Arosio, P. RNA Modulates HnRNPA1A Amyloid Formation Mediated by Biomolecular Condensates. Nat. Chem. 2024, 16 (7), 1052–1061. 10.1038/s41557-024-01467-3.

(48) Wadsworth, G. M.; Zahurancik, W. J.; Zeng, X.; Pullara, P.; Lai, L. B.; Sidharthan, V.; Pappu, R. V.; Gopalan, V.; Banerjee, P. R. RNAs Undergo Phase Transitions with Lower Critical Solution Temperatures. Nat. Chem. 2023, 15 (12), 1693–1704. 10.1038/s41557-023-01353-4.

(49) Mittag, T.; Parker, R. Multiple Modes of Protein–Protein Interactions Promote RNP Granule Assembly. Journal of Molecular Biology. Academic Press November 2, 2018, pp 4636–4649. 10.1016/j.jmb.2018.08.005.

(50) DeJesus-Hernandez, M.; Mackenzie, I. R.; Boeve, B. F.; Boxer, A. L.; Baker, M.; Rutherford, N. J.; Nicholson, A. M.; Finch, N. C. A.; Flynn, H.; Adamson, J.; Kouri, N.; Wojtas, A.; Sengdy, P.; Hsiung, G. Y. R.; Karydas, A.; Seeley, W. W.; Josephs, K. A.; Coppola, G.; Geschwind, D. H.; Wszolek, Z. K.; Feldman, H.; Knopman, D. S.; Petersen, R. C.; Miller, B. L.; Dickson, D. W.; Boylan, K. B.; Graff-Radford, N. R.; Rademakers, R. Expanded GGGGCC Hexanucleotide Repeat in Noncoding Region of C9ORF72 Causes Chromosome 9p-Linked FTD and ALS. Neuron 2011, 72 (2), 245–256. 10.1016/j.neuron.2011.09.011.

(51) Malik, I.; Kelley, C. P.; Wang, E. T.; Todd, P. K. Molecular Mechanisms Underlying Nucleotide Repeat Expansion Disorders. Nature Reviews Molecular Cell Biology. Nature Research September 1, 2021, pp 589–607. 10.1038/s41580-021-00382-6.

(52) Renton, A. E.; Majounie, E.; Waite, A.; Simón-Sánchez, J.; Rollinson, S.; Gibbs, J. R.; Schymick, J. C.; Laaksovirta, H.; van Swieten, J. C.; Myllykangas, L.; Kalimo, H.; Paetau, A.; Abramzon, Y.; Remes, A. M.; Kaganovich, A.; Scholz, S. W.; Duckworth, J.; Ding, J.; Harmer, D. W.; Hernandez, D. G.; Johnson, J. O.; Mok, K.; Ryten, M.; Trabzuni, D.; Guerreiro, R. J.; Orrell, R. W.; Neal, J.; Murray, A.; Pearson, J.; Jansen, I. E.; Sondervan, D.; Seelaar, H.; Blake, D.; Young, K.; Halliwell, N.; Callister, J. B.; Toulson, G.; Richardson, A.; Gerhard, A.; Snowden, J.; Mann, D.; Neary, D.; Nalls, M. A.; Peuralinna, T.; Jansson, L.; Isoviita, V. M.; Kaivorinne, A. L.; Hölttä-Vuori, M.; Ikonen, E.; Sulkava, R.; Benatar, M.; Wuu, J.; Chiò, A.; Restagno, G.; Borghero, G.; Sabatelli, M.; Heckerman, D.; Rogaeva, E.; Zinman, L.; Rothstein, J. D.; Sendtner, M.; Drepper, C.; Eichler, E. E.; Alkan, C.; Abdullaev, Z.; Pack, S. D.; Dutra, A.; Pak, E.; Hardy, J.; Singleton, A.; Williams, N. M.; Heutink, P.; Pickering-Brown, S.; Morris, H. R.; Tienari, P. J.; Traynor, B. J. A Hexanucleotide Repeat Expansion in C9ORF72 Is the Cause of Chromosome 9p21-Linked ALS-FTD. Neuron 2011, 72 (2), 257–268. 10.1016/j.neuron.2011.09.010.

(53) Ripin, N.; Parker, R. Are Stress Granules the RNA Analogs of Misfolded Protein Aggregates? 2022. 10.1261/rna.

(54) Budkina, K.; El Hage, K.; Clement, M. J.; Desforges, B.; Bouhss, A.; Joshi, V.; Maucuer, A.; Hamon, L.; Ovchinnikov, L. P.; Lyabin, D. N.; Pastre, D. YB-1 Unwinds MRNA Secondary Structures in Vitro and Negatively Regulates Stress Granule Assembly in HeLa Cells. Nucleic Acids Res. 2021, 49 (17), 10061–10081. 10.1093/nar/gkab748.

(55) Murthy, A. C.; Dignon, G. L.; Kan, Y.; Zerze, G. H.; Parekh, S. H.; Mittal, J.; Fawzi, N. L. Molecular Interactions Underlying Liquid−liquid Phase Separation of the FUS Low-Complexity Domain. Nat. Struct. Mol. Biol. 2019, 26 (7), 637–648. 10.1038/s41594-019-0250-x.

(56) Ren, C.-L.; Shan, Y.; Zhang, P.; Ding, H.-M.; Ma, Y.-Q. Uncovering the Molecular Mechanism for Dual Effect of ATP on Phase Separation in FUS Solution; 2022; Vol. 8. https://www.science.org.

(57) Ren, C.-L.; Shan, Y.; Zhang, P.; Ding, H.-M.; Ma, Y.-Q. Uncovering the Molecular Mechanism for Dual Effect of ATP on Phase Separation in FUS Solution; 2022; Vol. 8. https://www.science.org.

(58) Qamar, S.; Wang, G. Z.; Randle, S. J.; Ruggeri, F. S.; Varela, J. A.; Lin, J. Q.; Phillips, E. C.; Miyashita, A.; Williams, D.; Ströhl, F.; Meadows, W.; Ferry, R.; Dardov, V. J.; Tartaglia, G. G.; Farrer, L. A.; Kaminski Schierle, G. S.; Kaminski, C. F.; Holt, C. E.; Fraser, P. E.; Schmitt-Ulms, G.; Klenerman, D.; Knowles, T.; Vendruscolo, M.; St George-Hyslop, P. FUS Phase Separation Is Modulated by a Molecular Chaperone and Methylation of Arginine Cation-π Interactions. Cell 2018, 173 (3), 720–734.e15. 10.1016/j.cell.2018.03.056.

(59) Qamar, S.; Wang, G. Z.; Randle, S. J.; Ruggeri, F. S.; Varela, J. A.; Lin, J. Q.; Phillips, E. C.; Miyashita, A.; Williams, D.; Ströhl, F.; Meadows, W.; Ferry, R.; Dardov, V. J.; Tartaglia, G. G.; Farrer, L. A.; Kaminski Schierle, G. S.; Kaminski, C. F.; Holt, C. E.; Fraser, P. E.; Schmitt-Ulms, G.; Klenerman, D.; Knowles, T.; Vendruscolo, M.; St George-Hyslop, P. FUS Phase Separation Is Modulated by a Molecular Chaperone and Methylation of Arginine Cation-π Interactions. Cell 2018, 173 (3), 720–734.e15. 10.1016/j.cell.2018.03.056.

(60) Qamar, S.; Wang, G. Z.; Randle, S. J.; Ruggeri, F. S.; Varela, J. A.; Lin, J. Q.; Phillips, E. C.; Miyashita, A.; Williams, D.; Ströhl, F.; Meadows, W.; Ferry, R.; Dardov, V. J.; Tartaglia, G. G.; Farrer, L. A.; Kaminski Schierle, G. S.; Kaminski, C. F.; Holt, C. E.; Fraser, P. E.; Schmitt-Ulms, G.; Klenerman, D.; Knowles, T.; Vendruscolo, M.; St George-Hyslop, P. FUS Phase Separation Is Modulated by a Molecular Chaperone and Methylation of Arginine Cation-π Interactions. Cell 2018, 173 (3), 720–734.e15. 10.1016/j.cell.2018.03.056.

(61) Qamar, S.; Wang, G. Z.; Randle, S. J.; Ruggeri, F. S.; Varela, J. A.; Lin, J. Q.; Phillips, E. C.; Miyashita, A.; Williams, D.; Ströhl, F.; Meadows, W.; Ferry, R.; Dardov, V. J.; Tartaglia, G. G.; Farrer, L. A.; Kaminski Schierle, G. S.; Kaminski, C. F.; Holt, C. E.; Fraser, P. E.; Schmitt-Ulms, G.; Klenerman, D.; Knowles, T.; Vendruscolo, M.; St George-Hyslop, P. FUS Phase Separation Is Modulated by a Molecular Chaperone and Methylation of Arginine Cation-π Interactions. Cell 2018, 173 (3), 720–734.e15. 10.1016/j.cell.2018.03.056.

(62) Ahlers, J.; Adams, E. M.; Bader, V.; Pezzotti, S.; Winklhofer, K. F.; Tatzelt, J.; Havenith, M. The Key Role of Solvent in Condensation: Mapping Water in Liquid-Liquid Phase-Separated FUS. Biophys. J. 2021, 120 (7), 1266–1275. 10.1016/j.bpj.2021.01.019.

(63) Bentmann, E.; Neumann, M.; Tahirovic, S.; Rodde, R.; Dormann, D.; Haass, C. Requirements for Stress Granule Recruitment of Fused in Sarcoma (FUS) and TAR DNA-Binding Protein of 43 KDa (TDP-43). Journal of Biological Chemistry 2012, 287 (27), 23079–23094. 10.1074/jbc.M111.328757.

(64) Babinchak, W. M.; Haider, R.; Dumm, B. K.; Sarkar, P.; Surewicz, K.; Choi, J. K.; Surewicz, W. K. The Role of Liquid-Liquid Phase Separation in Aggregation of the TDP-43 Low-Complexity Domain. Journal of Biological Chemistry 2019, 294 (16), 6306–6317. 10.1074/jbc.RA118.007222.

(65) Russo, J.; Tavares, J. M.; Teixeira, P. I. C.; Telo Da Gama, M. M.; Sciortino, F. Reentrant Phase Diagram of Network Fluids. Phys. Rev. Lett. 2011, 106 (8), 1–4. 10.1103/PhysRevLett.106.085703.

(66) Zhang, F.; Skoda, M. W. A.; Jacobs, R. M. J.; Zorn, S.; Martin, R. A.; Martin, C. M.; Clark, G. F.; Weggler, S.; Hildebrandt, A.; Kohlbacher, O.; Schreiber, F. Reentrant Condensation of Proteins in Solution Induced by Multivalent Counterions. Phys. Rev. Lett. 2008, 101 (14), 3–6. 10.1103/PhysRevLett.101.148101.

(67) Milin, A. N.; Deniz, A. A. Reentrant Phase Transitions and Non-Equilibrium Dynamics in Membraneless Organelles. Biochemistry 2018, 57 (17), 2470–2477. 10.1021/acs.biochem.8b00001.

(68) Levy, Y.; Hazra, M. K. Biophysics of Phase Separation of Disordered Proteins Is Governed by Balance between Short- And Long-Range Interactions. Journal of Physical Chemistry B 2021, 125 (9), 2202–2211. 10.1021/acs.jpcb.0c09975.

(69) Hazra, M. K.; Levy, Y. Charge Pattern Affects the Structure and Dynamics of Polyampholyte Condensates. Physical Chemistry Chemical Physics 2020. 10.1039/d0cp02764b.

(70) Givaty, O.; Levy, Y. Protein Sliding along DNA: Dynamics and Structural Characterization. J. Mol. Biol. 2009, 385 (4), 1087–1097. 10.1016/j.jmb.2008.11.016.

(71) Dignon, G. L.; Zheng, W.; Kim, Y. C.; Best, R. B.; Mittal, J. Sequence Determinants of Protein Phase Behavior from a Coarse-Grained Model. PLoS Comput. Biol. 2018, 14 (1). 10.1371/journal.pcbi.1005941.

(72) Kapoor, U.; Kim, Y. C.; Mittal, J. Coarse-Grained Models to Study Protein-DNA Interactions and Liquid-Liquid Phase Separation. J. Chem. Theory Comput. 2023. 10.1021/acs.jctc.3c00525.

