## Supporting Information for "Dual Effect of RNA-like Polyelectrolytes on Stability and Dynamics of Biomolecular Condensates: A Tale of Competitive Short and Long-range Interactions"

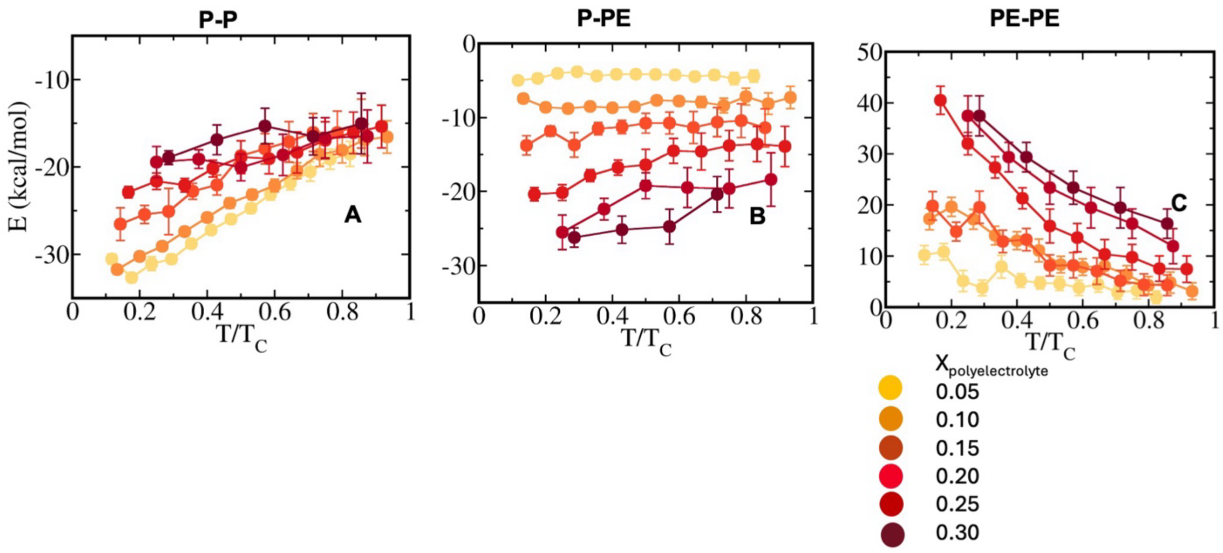


Fig S1. Dissection of interaction energy into self and cross components in peptide–polyelectrolyte condensates for sequence pair f = 0.0 and $(U)_{20}$. Average interaction energy per chain originating from (A) peptide–peptide (P–P), (B) peptide-polyelectrolyte (P–PE), and (C) polyelectrolyte-polyelectrolyte (PE–PE) components has been shown as a function of temperature scaled to criticality at various polyelectrolyte concentrations. The colour code indicates polyelectrolyte mole fractions ($X_{Polyelectrolyte}$), ranging from yellow (low concentration) to dark red (high concentration).

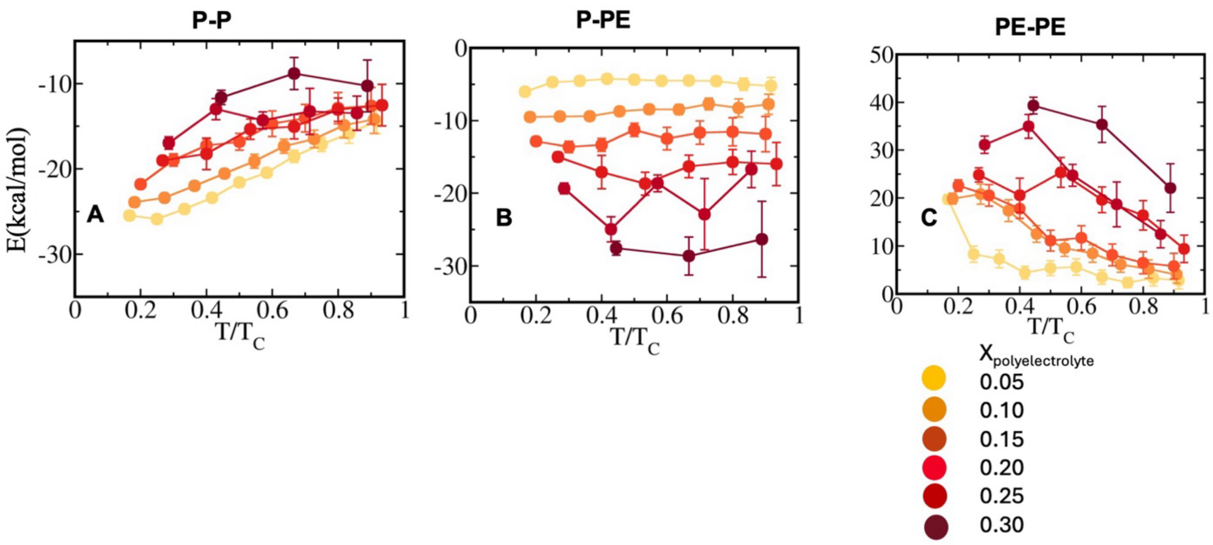


Fig S2. Dissection of interaction energy into self and cross components in peptide–polyelectrolyte condensates for sequence pairs f = 0.1 and $(U)_{20}$.Average interaction energy per chain originating from (A) peptide–peptide (P–P), (B) peptide-polyelectrolyte (P–PE), and (C) polyelectrolyte-polyelectrolyte (PE–PE) components has been shown as a function of temperature scaled to criticality at various polyelectrolyte concentrations. The colour code indicates polyelectrolyte mole fractions ($X_{Polyelectrolyte}$), ranging from yellow (low concentration) to dark red (high concentration).

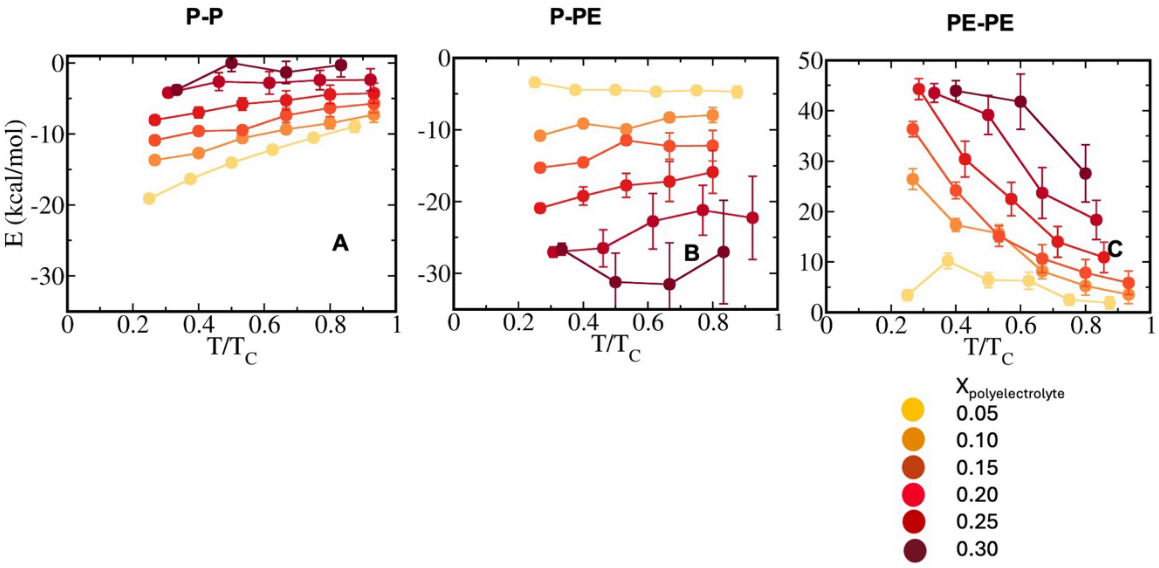


Fig S3. Dissection of interaction energy into self and cross components in peptide–polyelectrolyte condensates for sequence pairs f = 0.3 and $(U)_{20}$. Average interaction energy per chain originating from (A) peptide–peptide (P–P), (B) peptide-polyelectrolyte (P–PE), and (C) polyelectrolyte-polyelectrolyte (PE–PE) components has been shown as a function of temperature scaled to criticality at various polyelectrolyte concentrations. The colour code indicates polyelectrolyte mole fractions ($X_{Polyelectrolyte}$), ranging from yellow (low concentration) to dark red (high concentration).

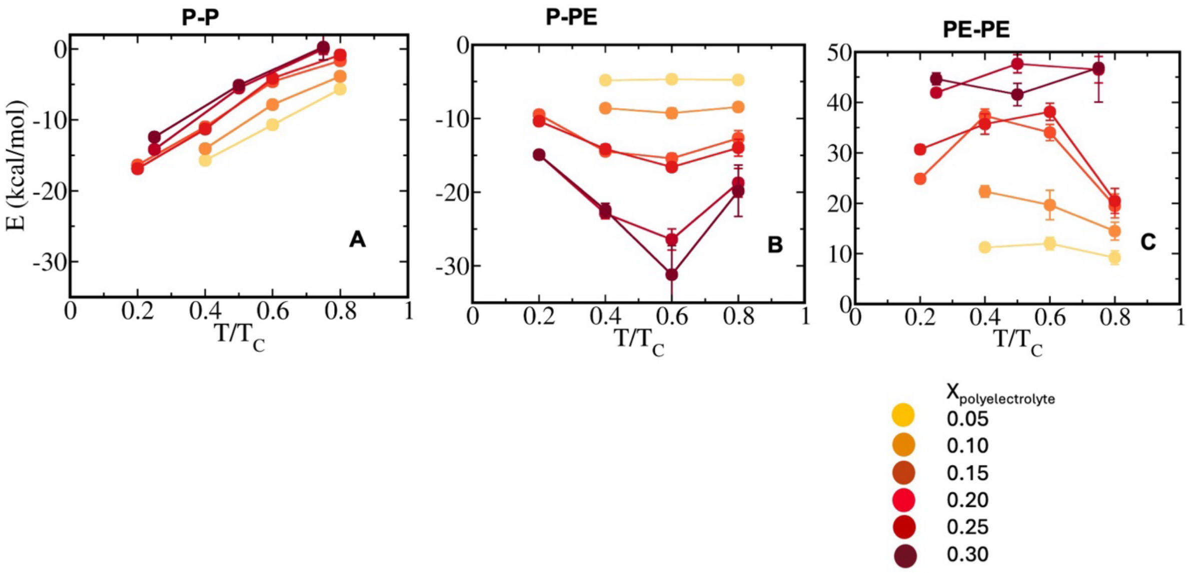


Fig S4. Dissection of interaction energy into self and cross components in peptide–polyelectrolyte condensates for sequence pairs f = 0.6 and $(U)_{20}$. Average interaction energy per chain originating from (A) peptide–peptide (P–P), (B) peptide-polyelectrolyte (P–PE), and (C) polyelectrolyte-polyelectrolyte (PE–PE) components has been shown as a function of temperature scaled to criticality at various polyelectrolyte concentrations. The colour code indicates polyelectrolyte mole fractions ($X_{Polyelectrolyte}$), ranging from yellow (low concentration) to dark red (high concentration).

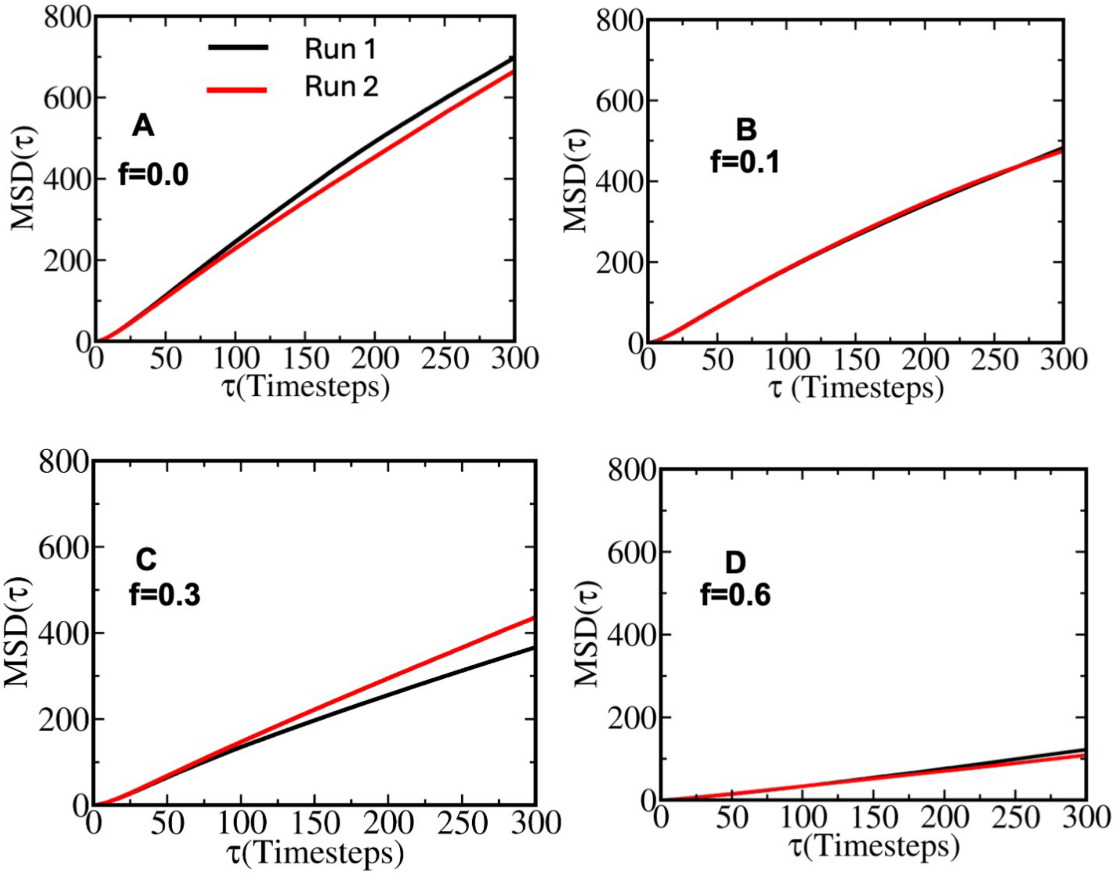


Fig S5. Convergence of mean squared displacement (MSD) of peptides in condensate phase from independent simulation runs for sequence hydrophobicity (A) f = 0.0 (B) f=0.1 (C) f=0.3 and (D) f=0.6 at T/TC=0.6 and polyelectrolyte mixing fraction $X_{Polyelectrolyte} = 0.15$. The close overlap between the MSD profiles confirms the reproducibility of droplet-phase dynamics across independent simulations

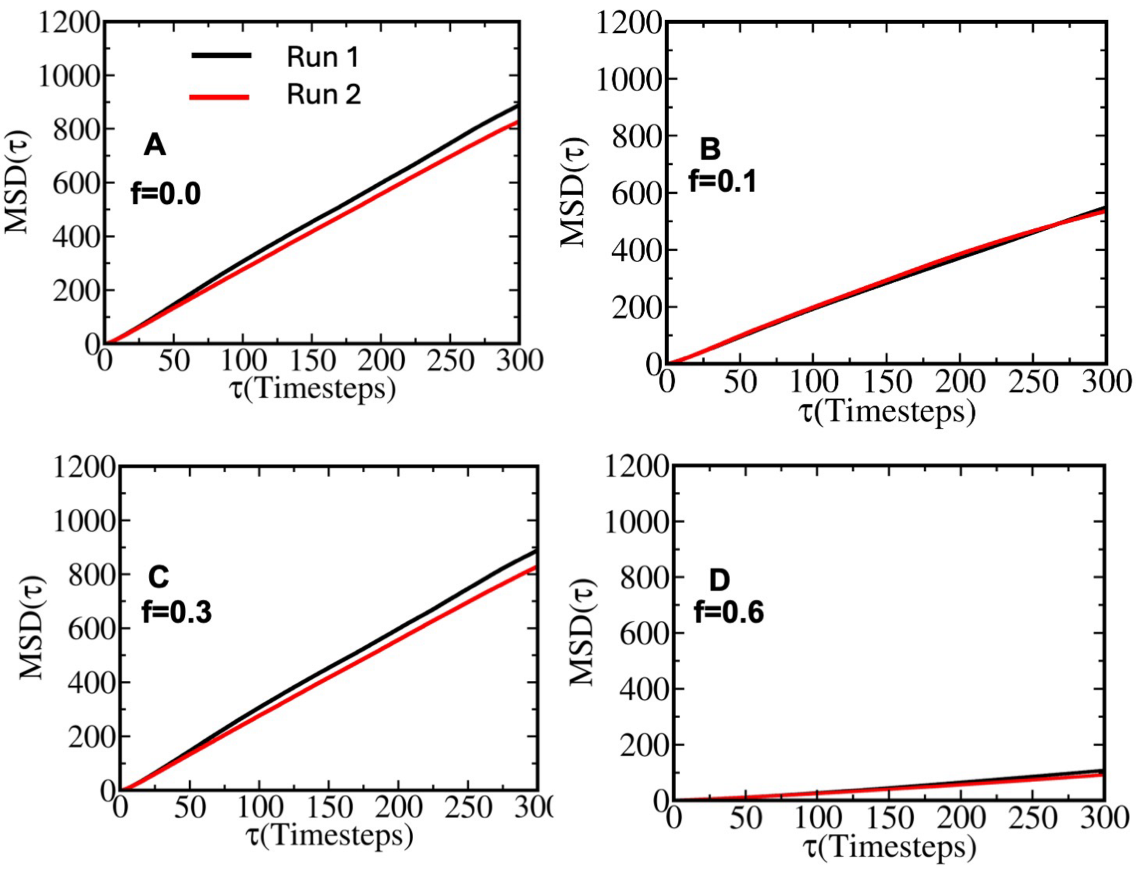


Fig S6. Convergence of mean squared displacement (MSD) of polyelectrolytes in condensate phase from independent simulation runs for sequence hydrophobicity (A) f = 0.0 (B) f=0.1 (C) f=0.3 and (D) f=0.6 at T/TC=0.6 and polyelectrolyte mixing fraction $X_{Polyelectrolyte} = 0.15$. The close overlap between the MSD profiles confirms the reproducibility of droplet-phase dynamics across independent simulations.

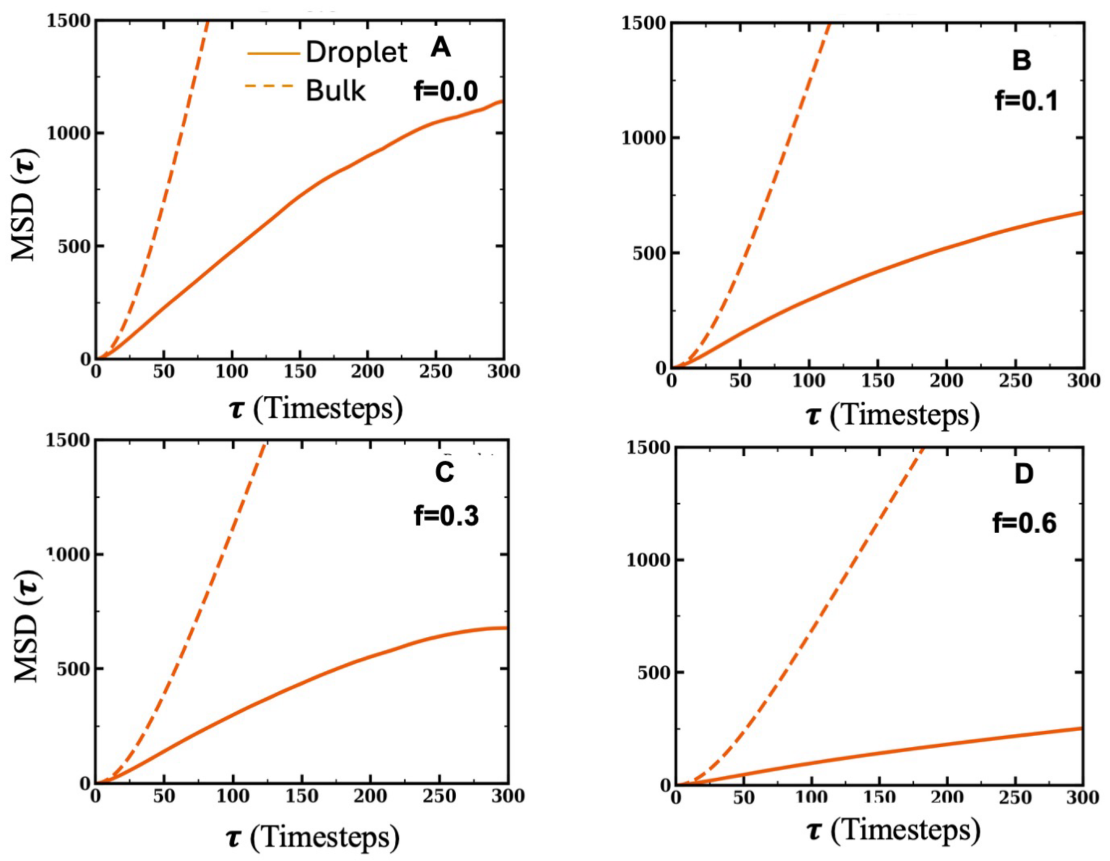


Fig S7. Comparative mean squared displacement (MSD) of peptides in droplet and bulk phases at fixed distance from criticality for sequences with hydrophobicity content (A) f = 0.0, (B) f = 0.1, (C) f = 0.3, and (D) f = 0.6, at a fixed polyelectrolyte concentration ($X_{Polyelectrolyte} = 0.15$). Solid lines represent MSD measured inside the droplet (condensate) phase, while dashed lines correspond to MSD in the bulk phase.

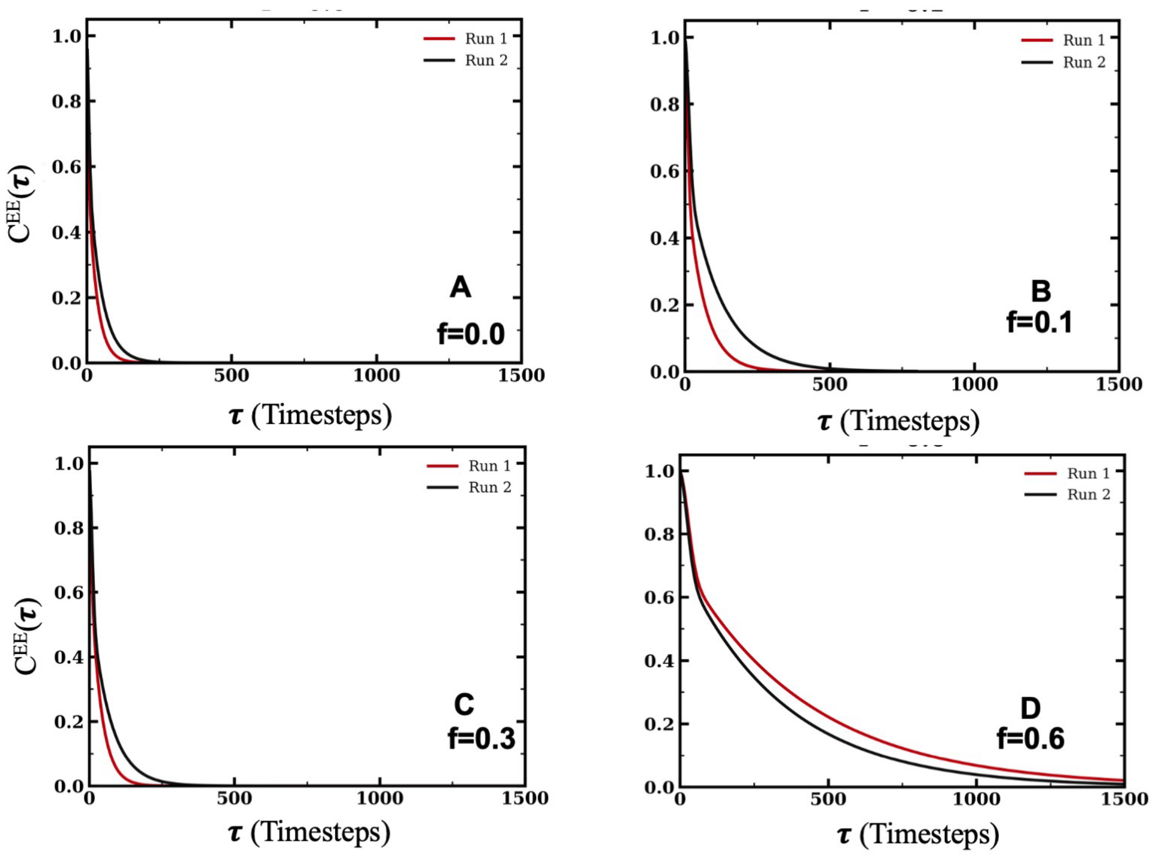


Fig S8. Convergence of end-to-end distance time correlation function of peptides in droplet phase at $X_{Polyelectrolyte} = 0.15$ and T/Tc=0.6 obtained from two independent simulation runs for condensate forming sequences (A) f = 0.0, (B) f = 0.1, (C) f = 0.3, and (D) f = 0.6 as representative examples.

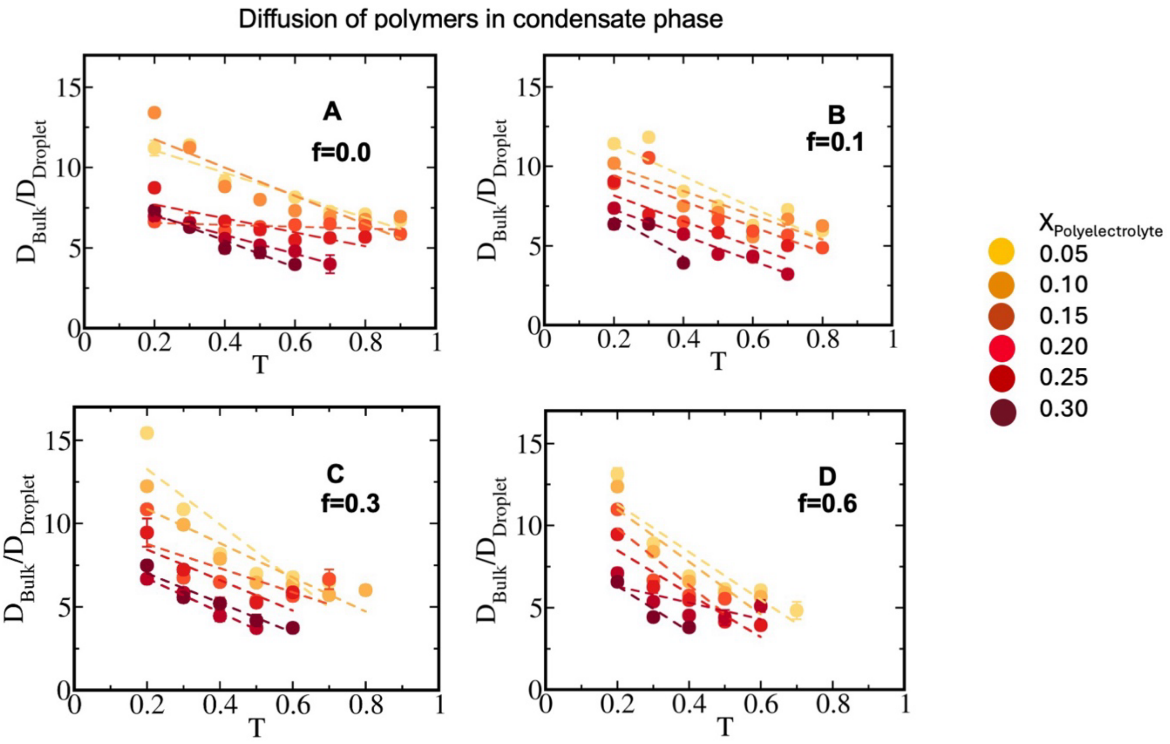


Fig S9. Droplet-to-bulk diffusivity contrast across hydrophobicity and RNA-like polyelectrolyte concentration. The ratio $D_{bulk}/D_{droplet}$ has been shown as a function of absolute temperature for different hydrophobicity strengths: (A) f = 0.0, (B) f = 0.1, (C) f = 0.3, and (D) f = 0.6, across polyelectrolyte concentrations. Colour codes denote increase in polyelectrolyte mixing fraction, ranging from yellow (low concentration) to dark red (high concentration).

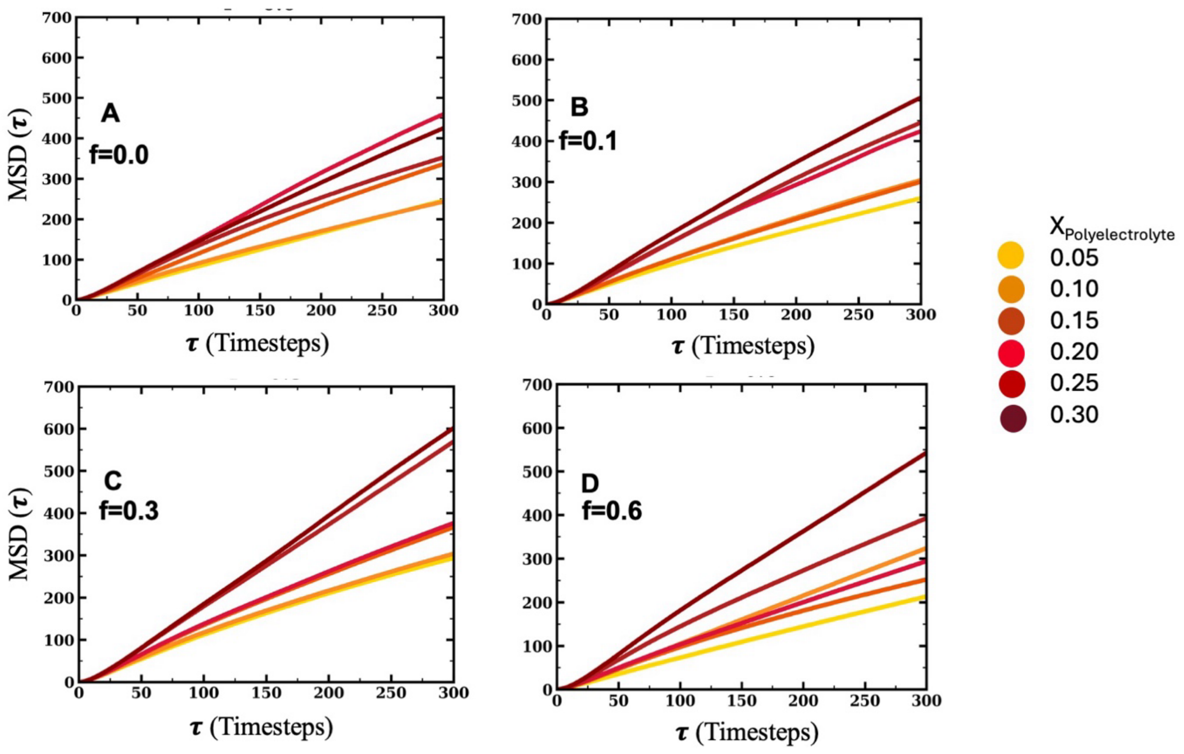


Fig S10. Average mean squared displacement (MSD) of peptides in condensates along polyelectrolyte mixing fraction at fixed temperature (T = 0.3) for different hydrophobicity content of the sequences forming condensates: (A) f = 0.0, (B) f = 0.1, (C) f = 0.3, and (D) f = 0.6, Color codes denote polyelectrolyte mixing fraction ($X_{Polyelectrolyte}$) , ranging from yellow (low concentration) to dark-red (high concentration).

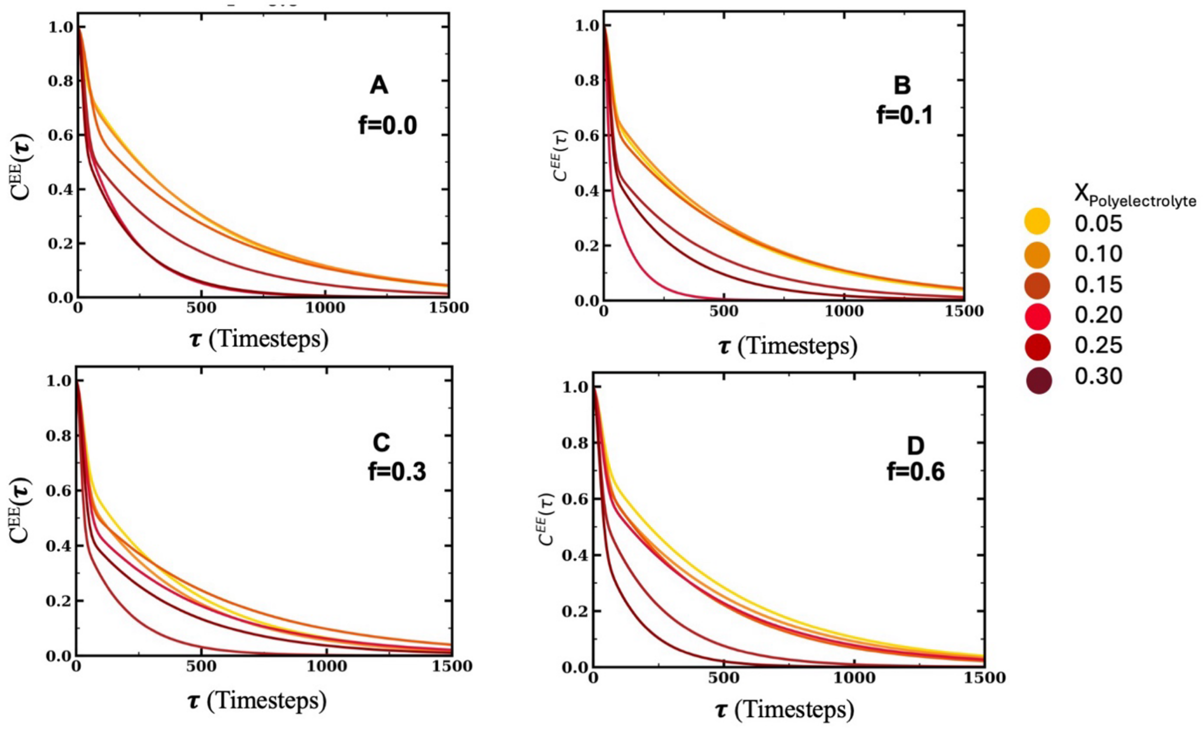


Fig S11. Average end-to-end distance time correlation function of peptides in condensates along polyelectrolyte mixing fraction at fixed temperature (T = 0.3) for sequences having different hydrophobicity content forming condensates: (A) f = 0.0, (B) f = 0.1, (C) f = 0.3, and (D) f = 0.6, Color codes denote polyelectrolyte mixing fraction ($X_{Polyelectrolyte}$) , ranging from yellow (low concentration) to dark-red (high concentration).

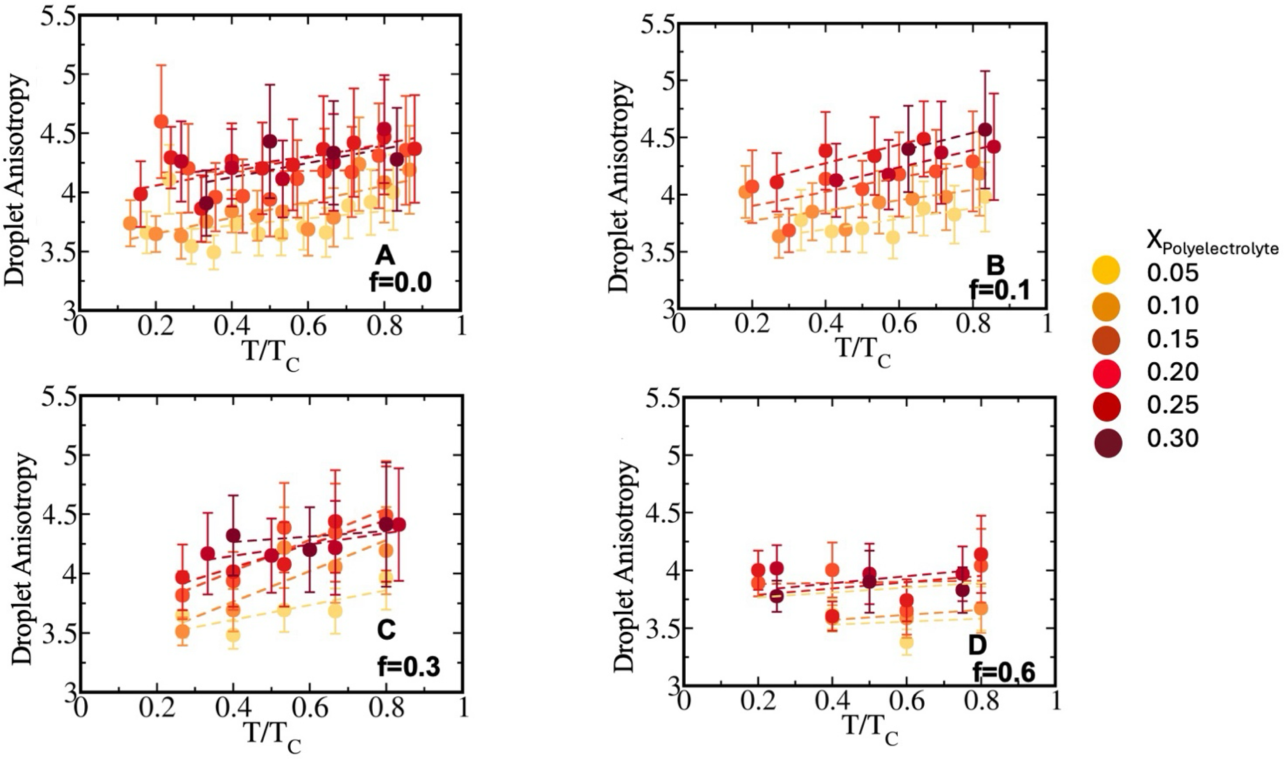


Fig S12. Droplet shape anisotropy as a function of temperature scaled to criticality. The shape anisotropy of the largest condensate cluster is shown as a function temperature for different hydrophobicity strengths of the designed sequences: (A) f = 0.0, (B) f = 0.1, (C) f = 0.3, and (D) f = 0.6, at various polyelectrolyte concentrations ranging from $X_{Polyelectrolyte} = 0.05 - 0.30$ . Colour codes represent the polyelectrolyte concentration, ranging from yellow (low concentration) to dark red (high concentration).

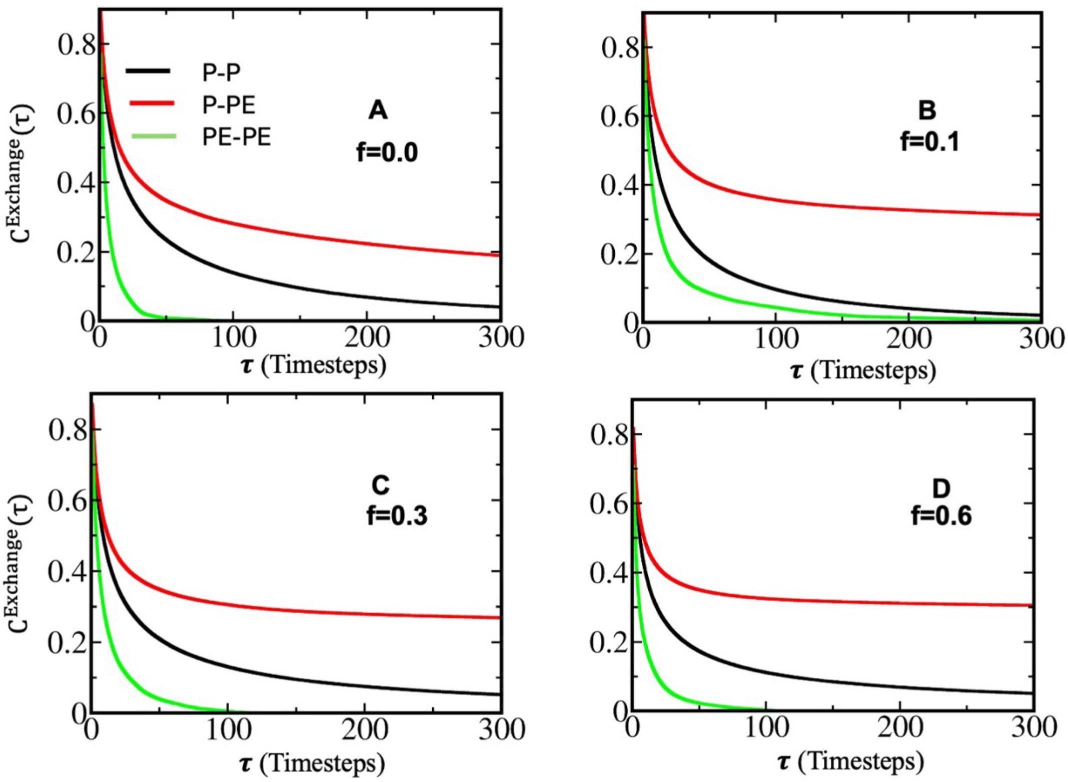


Fig S13. Neighbour Contact exchange time correlation function in condensate phase of sequence pairs (A) f = 0.0 and $(U)_{20}$, (B) f = 0.1 and $(U)_{20}$, (C) f = 0.3 and $(U)_{20}$, and (D) f = 0.6 and $(U)_{20.}$ Each panel showcase peptide-peptide contact time correlation (P-P) in black, peptide-polyelectrolyte contact time correlations (P-PE) in red and polyelectrolyte-polyelectrolyte (PE-PE) contact time correlations with green. While P-P and PE-PE neighbours are fairly exchanging, P-PE contacts are sticky.
